# Heat Shock Protein 27 Immune Complex Upregulates LDLR Expression Thereby Reducing Plasma Cholesterol and Atherogenesis

**DOI:** 10.1101/2020.05.21.102350

**Authors:** Yong-Xiang Chen, Chunhuai Shi, Jingti Deng, Catherine Diao, Nadia Maarouf, Matthew Rosin, Vipul Shrivastava, Angie A. Hu, Sonya Bharadwa, Ayinuer Adijiang, Annegret Ulke-Lemee, Brenig Gwilym, Alexandria Hellmich, Christopher Malozzi, Zarah Batulan, Jonathan L. E. Dean, F. Daniel Ramirez, Jingwen Liu, William T. Gerthoffer, Edward R. O’Brien

## Abstract

Elevated Heat Shock Protein 27 levels predict relative freedom from cardiovascular events. In ApoE^-/-^ mice HSP27 over-expression or twice daily subcutaneous injections reduce blood and plaque cholesterol levels, inflammation and atherogenesis. While natural antibodies to HSP27 are present in human blood their role is unknown. Here, we show that blood levels of both HSP27 and anti-HSP27 IgG antibodies are elevated in healthy controls compared to patients with cardiovascular disease. ApoE^-/-^ mice vaccinated with recombinant HSP25 (murine ortholog) develop elevated anti-HSP25 IgG antibodies and reduced levels of cholesterol, inflammation and atherosclerosis. The effects on cholesterol metabolism were divergent: increased hepatic LDLR expression and reduced plasma PCSK9 levels. *In vitro*, a polyclonal anti-HSP27 IgG antibody combined with rHSP27 to upregulate hepatocyte LDLR expression via an NF-kB-dependent pathway that is independent of SREBP2 expression and intracellular cholesterol levels. HSP27 immunotherapy represents a novel means of lowering not only cholesterol but also PCSK9.

---

Small heat shock proteins, such as Heat Shock Protein 27 (HSP27), are intracellular chaperones that promote the proper reassembly of misfolded proteins and act as mediators of extracellular cellular signaling^1^. HSP27 effectively preserves cellular homeostasis under various conditions of degenerative or inflammatory stress – including those common to the pathogenesis of atherosclerosis^2^. There is overwhelming genetic, epidemiological and clinical evidence that irrefutably establishes low density lipoprotein cholesterol (LDL-C) as causal for atherosclerosis^3, 4^. Experiments from our group^5^ and four others using proteomic discovery approaches^6–9^ show that serum HSP27 levels decline as human atherosclerosis develops, with its tissue abundance inversely corelated with the degree of coronary artery plaque burden. In atherosclerosis-prone Apolipoprotein E null (*ApoE^-/-^*) mice augmentation of extracellular HSP27 levels via constitutive over-expression, transplantation of bone marrow from mice that over-express HSP27, twice-daily subcutaneous administration of recombinant HSP27 (rHSP27; 100 μg) or estrogenic therapy post-ovariectomy (that augment HSP27 blood levels) reduce both plasma and plaque cholesterol content, resulting in the formation of more stable plaques that are less inflamed^10–13^. Clinically, elevated HSP27 blood levels are associated with a lower 5-year risk of myocardial infarction, stroke or cardiovascular death^11^. Interestingly, natural antibodies to HSP27 (AAbs) are detectable in the blood, yet their biological significance is unclear^14, 15^.

In this study we sought to address three questions. First, what is the correlation between blood HSP27 and AAb abundance in human cardiovascular disease (CVD) patients compared to healthy control subjects (CON)? Second, does augmenting levels of antibodies to HSP25 (the murine ortholog of human HSP27) via vaccination attenuate atherogenesis in *ApoE^-/-^* mice?

Third, what are the potential mechanisms by which the HSP27 immune complex (IC) alters the two key drivers of atherosclerosis: cholesterol and inflammation? Our results indicate that blood levels of HSP27 and AAbs are higher in health compared to CVD. rHSP25 vaccination boosts anti-HSP25 antibodies and is associated with reductions in plasma cholesterol as well as proprotein convertase subtilisin / kexin type 9 (PCSK9) levels^16^, a key negative regulator of low density lipoprotein receptor (LDLR) recycling. These anti-atherogenesis effects are associated with reduced plaque and hepatic inflammation. Of novel importance, the HSP27 IC markedly upregulates LDLR expression independent of intracellular cholesterol levels and is reliant on activation of the NF-*κ*B pathway. Taken together, the discovery of HSP27-mediated upregulation of LDLR heralds a unique opportunity to develop HSP27 immuno-therapeutics for the treatment of atherosclerosis and hypercholesterolemia.

## Results

### Anti-HSP27 IgG antibodies are more abundant in health *vs*. CVD

To assess HSP27 and AAb levels, 80 subjects with cardiovascular disease (CVD) and 58 controls (CON) were recruited as part of a National Institutes of Health (NIH) sponsored study designed to look for novel cardiovascular biomarkers in a medically under-serviced population (**Supplemental Table 1**). Similar to our previous report^11^, CVD was associated with a lower HSP27 blood concentration compared to CON (586.2 ± 25.9 *vs.* 724.9 ± 29.8 pg/mL, p<0.0006). This difference persisted after adjusting for age, sex, race, BMI, diabetes mellitus, and active smoking status (adjusted difference: −117.1 pg/mL, 95% CI-221.4 to −12.8, p=0.028; **Fig. 1A**). Diabetes mellitus was associated with lower HSP27 levels in this model (adjusted difference: −127.4 pg/mL, 95% CI-232.7 to −22.2; p=0.0181) whereas the remaining parameters were not significant, including sex (p=0.0929) and ethnicity (p=0.7417 and 0.6632 for African American and Other, respectively).

A new insight is that CVD was associated with lower IgG AAb levels (31.3 ± 2.6 *vs*. 54.7 ± 4.1 arbitrary units [a.u.]: crude analysis: p<0.0001; adjusted difference: −21.8 a.u., 95% CI-33.6 to −10.0, p=0.0004; **Fig. 1B**). The association of CVD with IgM AAb levels was not significant after adjusting for prespecified clinical variables (crude difference: 0.161 ± 0.112 *vs.* 0.214 ± 0.144 a.u., p=0.022; adjusted difference: −0.004 a.u., 95% CI-0.060 to 0.052; p=0.889). African American ethnicity was associated with higher IgG AAb levels relative to Caucasians (adjusted difference: ±15.5 a.u., 95% CI 5.4 to 25.6; p=0.003). No other parameters were associated with significant differences in IgG AAbs, including sex (p=0.07). Finally, the ratio of IgG AAb to HSP27 was higher in CON *vs*. CVD patients (0.083 ± 0.007 *vs*. 0.059 ± 0.005, p=0.007), thereby suggesting that binding of more IgG AAb to HSP27 may facilitate its anti-atherosclerosis effects (**Fig. 1C**). One individual with CVD had an IgG AAb:HSP27 ratio >5.0 standard deviations (SDs) above the CVD group mean. Excluding this outlier identified a weakly positive association between IgG AAbs and HSP27 (r=0.174, p=0.042) but did not appreciably change the above results.

### rHSP25 Treatment Attenuates Atherogenesis in *ApoE^-/-^* mice

With our clinical data showing elevated levels of both HSP27 and AAbs in health *vs*. CVD, we then tested the hypothesis that enhancing serum levels of these AAbs might be atheroprotective. Atherosclerosis-prone *ApoE^-/-^* mice were started on a high fat diet (HFD) for 2 weeks before receiving four weekly subcutaneous injections of 100 µg of recombinant HSP25 (rHSP25) mixed with the adjuvant Alum (2% aluminum hydroxide; 3:1 v/v, **Fig. 2A**). Control mice were treated with rC1, the recombinant C-terminal of HSP27 that is biologically inactive (100 µg mixed with Alum). The use of rC1 is particularly relevant as a control, because rC1 and rHSP25 are both generated in *E. coli*. While the endotoxin contamination concentrations are lower than 2 units/mg of each recombinant protein, any potential confounding endotoxin effects are counterbalanced by comparing the active and control treatment groups. Separately, we validated rC1 as a control treatment, noting that [PBS + Alum] and [rC1 + Alum] were equally ineffective in attenuating atherogenesis and lowering cholesterol levels (**Supplemental Fig. 1A-1B**), even though vaccination with [rC1 + Alum] was similar to [rHSP25 + Alum] in generating AAbs, while [PBS + Alum] was not (**Fig. 2B**).

**Fig. 1.**
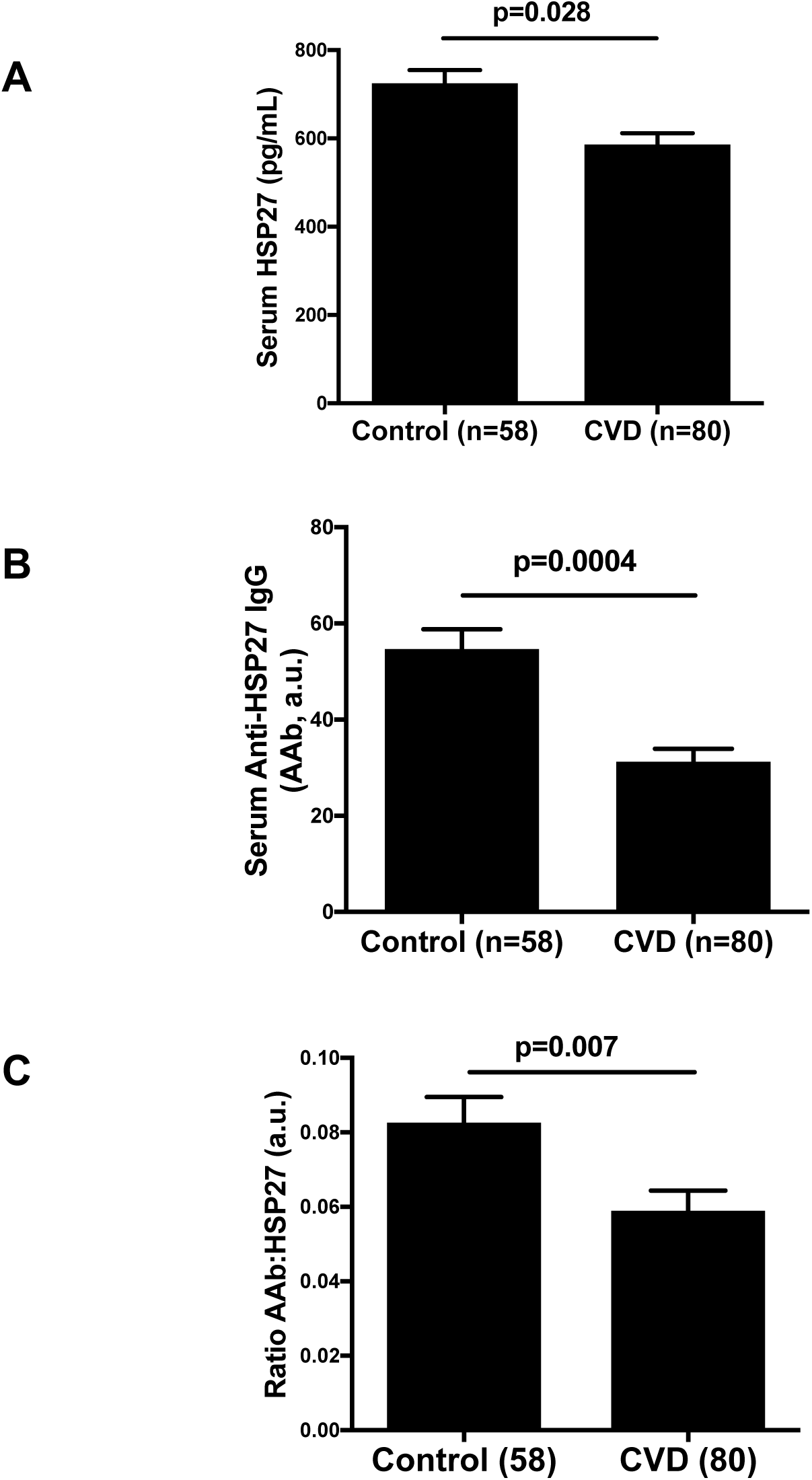
Serum HSP27 and anti-HSP27 autoantibody (AAb) levels in healthy human controls (CON) and cardiovascular disease (CVD) patients. All of the following parameters are higher in CON vs. CVD: A) Serum HSP27 levels B)Serum AAb levels C)The ratio of the AAb to HSP27

**Fig. 2.**
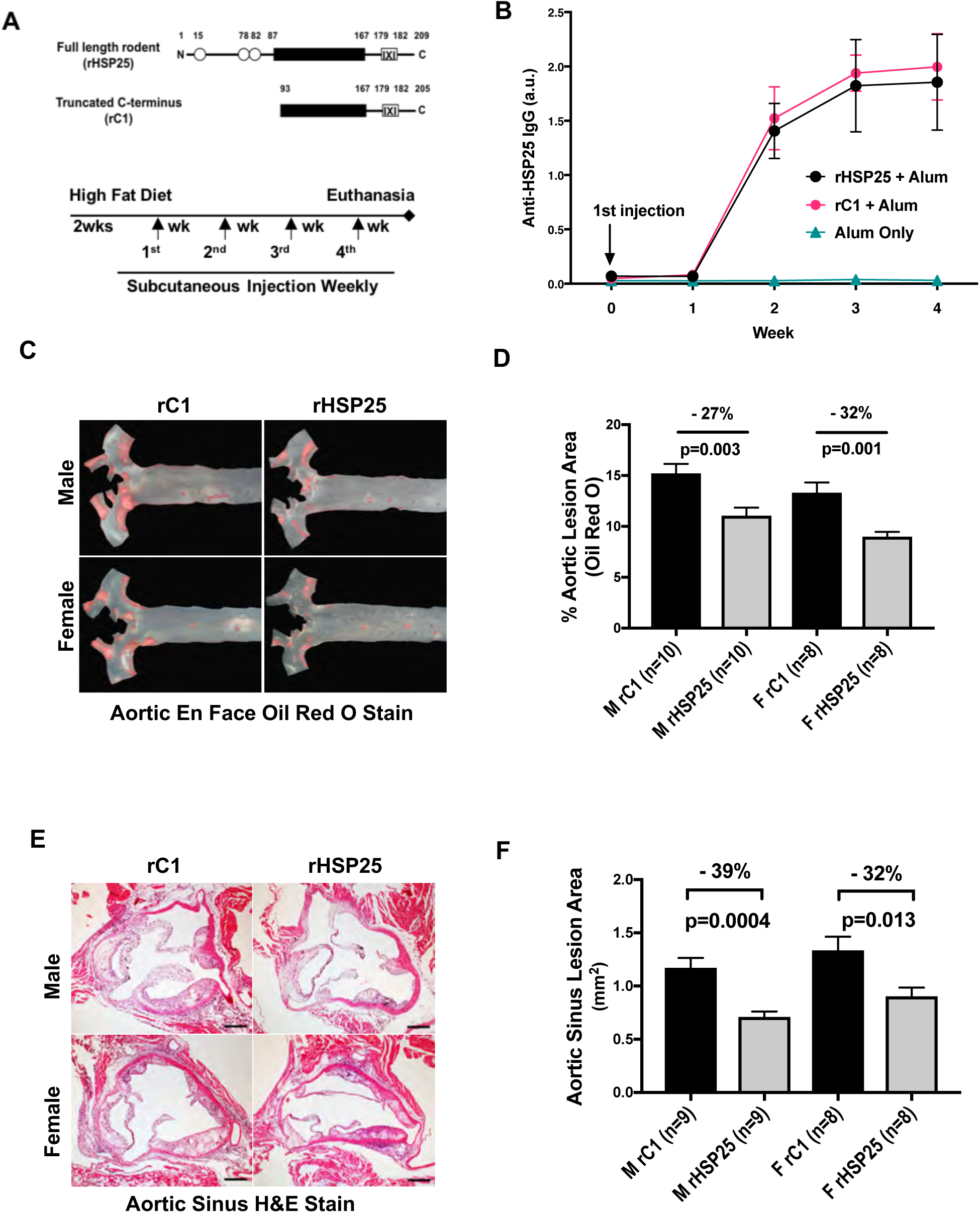
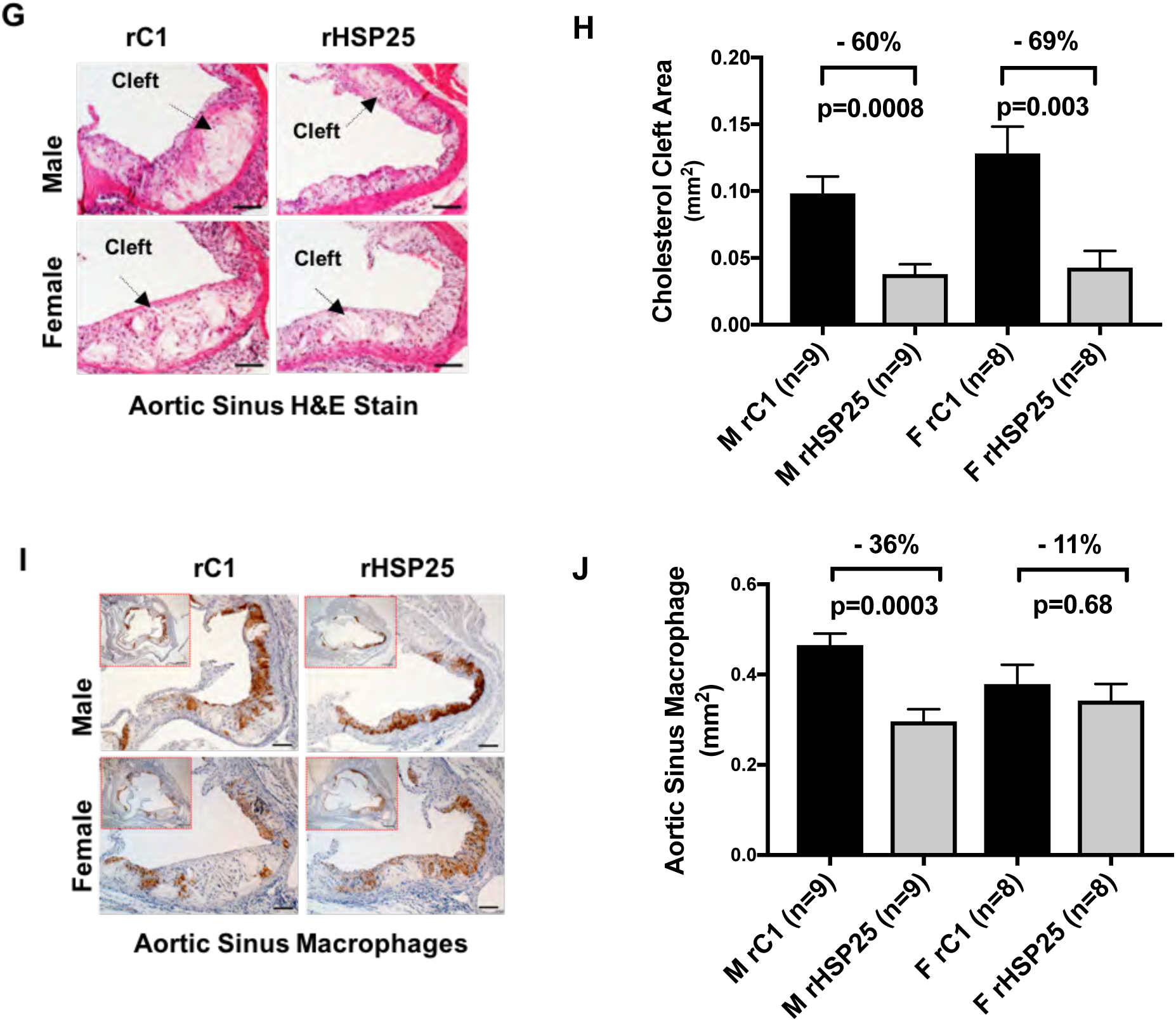
rHSP25 immunotherapy strategy attenuates atherosclerosis in *ApoE^-/-^* mice. A) Schematic representation of rHSP25 (mouse ortholog of HSP27) and the biologically inactive truncated C-terminal of HSP27 (rC1 control) proteins. The black box denotes the alpha beta-crystallin domain that is important for protein oligomerization. The IXI box at the C-terminus is a flexible domain involved in the formation of multiple inter-subunit interactions. The timeline for the murine treatment experiments is shown below, highlighting the duration of the high fat diet, and the weekly subcutaneous injections of rHSP25 or rC1 with the adjuvant alum. B) Time course for the *in vivo* generation of IgG antibodies in *ApoE^-/-^* mice after weekly treatment with rHSP25 and rC1 but not PBS. (a.u. = arbitrary units for OD @ 450 nm). C) and D) Aortic *en face* lesion area visualized with Oil Red O denoting neutral lipid deposits. The aortic lesion area was reduced in male and female mice vaccinated with rHSP25 *vs.* rC1. E) and F) Aortic sinus cross-sections stained with hematoxylin and eosin (H&E). Atherosclerotic lesion areas were reduced in male and female mice vaccinated with rHSP25 *vs.* rC1. Scale bar = 0.5 mm. G) and H) rHSP25 vaccination reduced the plaque cholesterol cleft content in male and female mice compared to rC1 control treatment. Scale bar = 0.2 mm. I) and J) Mouse plaque M*Φ* content was reduced by rHSP25 vaccination in male mice but was not significantly reduced in female mice. The brown color reaction product identifies M*Φ* immunolabeled with an anti-Mac-2 antibody. Scale bar = 0.2 mm for larger images, and 0.5 mm for insert images.

Atherosclerosis, as reflected by the *en face* aortic area of oil red O staining (indicative of intracellular lipid deposits), was reduced by 27% with rHSP25 treatment in male (15.2% ± 0.9% *vs.* 11.1% ± 8.0%, n=10/group; p=0.003) and 32% in female mice (13.3% ± 1.0% *vs*. 9.0% ± 0.5%, n=8/group; p=0.001; **Fig. 2C, 2D**). Similarly, the amount of atherosclerotic plaque quantified from aortic sinus cross-sections (**Fig. 2E, 2F**) was reduced 39% in male (1.17 ± 0.09 *vs*. 0.71 ± 0.05, n=9/group; p=0.0004) and 32% in female mice (1.34 ± 0.13 *vs*. 0.91 ± 0.08, n=8/group; p=0.013). These changes in plaque burden were accompanied by alterations in the content of the plaques. The cholesterol cleft area of the plaques (**Fig. 2G, 2H**) was reduced by 60% in male (0.10 ± 0.01 *vs*. 0.04 ± 0.01, n= 9/group; p=0.0008) and 69% in female mice (0.13 ± 0.02 *vs*. 0.04 ± 0.01, n=8/group; p=0.0027). However, there were sex differences with respect to the reduction in plaque macrophage (M*Φ*) content in response to treatment (**Fig. 2I, 2J**). rHSP25 treatment reduced plaque M*Φ* content by 36% in male (0.47 ± 0.03 *vs.* 0.30 ± 0.03, p=9/group; p=0.0003) but not female mice (a 11% reduction was not significant: 0.38 ± 0.04 *vs*. 0.34 ± 0.04, n=8/group; p=0.68). Overall, there was a strong relationship between lesion area and either cholesterol cleft area (r^2^=0.626; p<0.001) or M*Φ* area (r^2^=0.274; p=0.002).

Of note, rHSP25 vaccination of LDLR^-/-^ mice was ineffective in altering total plasma cholesterol (**Supplemental Fig. 2A**) – suggesting that the LDLR is necessary for HSP25-mediated cholesterol lowering. Vaccination with rHSP25 did not reduce plasma cholesterol levels in *ApoE^-/-^* mice that had HSP25 knocked out (*ApoE^-/-^HSP25^-/-^*), despite a robust antibody response to both rHSP25 and rC1 (**Supplemental Fig. 2B**). Therefore, it appears that at least basal levels of endogenous HSP25 are required for AAbs to lower cholesterol levels.

### rHSP25 Vaccination Reduces Hepatic Inflammation

In that hepatic inflammation may reflect vessel wall inflammation^17^ we assessed hepatic lipid-laden M*Φ* content by immunolabeling with an anti-Mac-2 antibody and performing concomitant oil red O staining. Compared to rC1, rHSP25 vaccination dramatically decreased hepatic M*Φ* content in male (70%, 10.6 ± 0.4 *vs*. 3.2 ± 0.4, n=10/group; p<0.0001) and female mice (60%, 11.3 ± 0.9 *vs.* 4.5 ± 0.3, n=6/group; p<0.0001) (**Fig. 3A, 3B**). In addition, the expression of inflammatory M*Φ* markers, as well as cytokines (as determined using qPCR) was lower in liver tissue from mice treated with rHSP25 *vs*. rC1 (n=3 F + n=3 M, per group). For male mice (**Fig. 3C**) the changes were as follows: CD68 (−63%, p=0.006), IL-1*β* (−15%, p=0.06), MCP1 (−41%, p=0.02) and TNF1*α* (−55%, p=0.003), with no change in IL-10 expression (p=0.59). Similarly, for female mice (**Fig. 3D**) the changes were as follows: CD68 (−48%, p=0.008), IL-1*β* (−57%, p=0.004), MCP1 (−55%, p=0.01) and TNF1*α* (−52%, p=0.005), with no change in IL-10 expression (p=0.98).

**Fig. 3.**
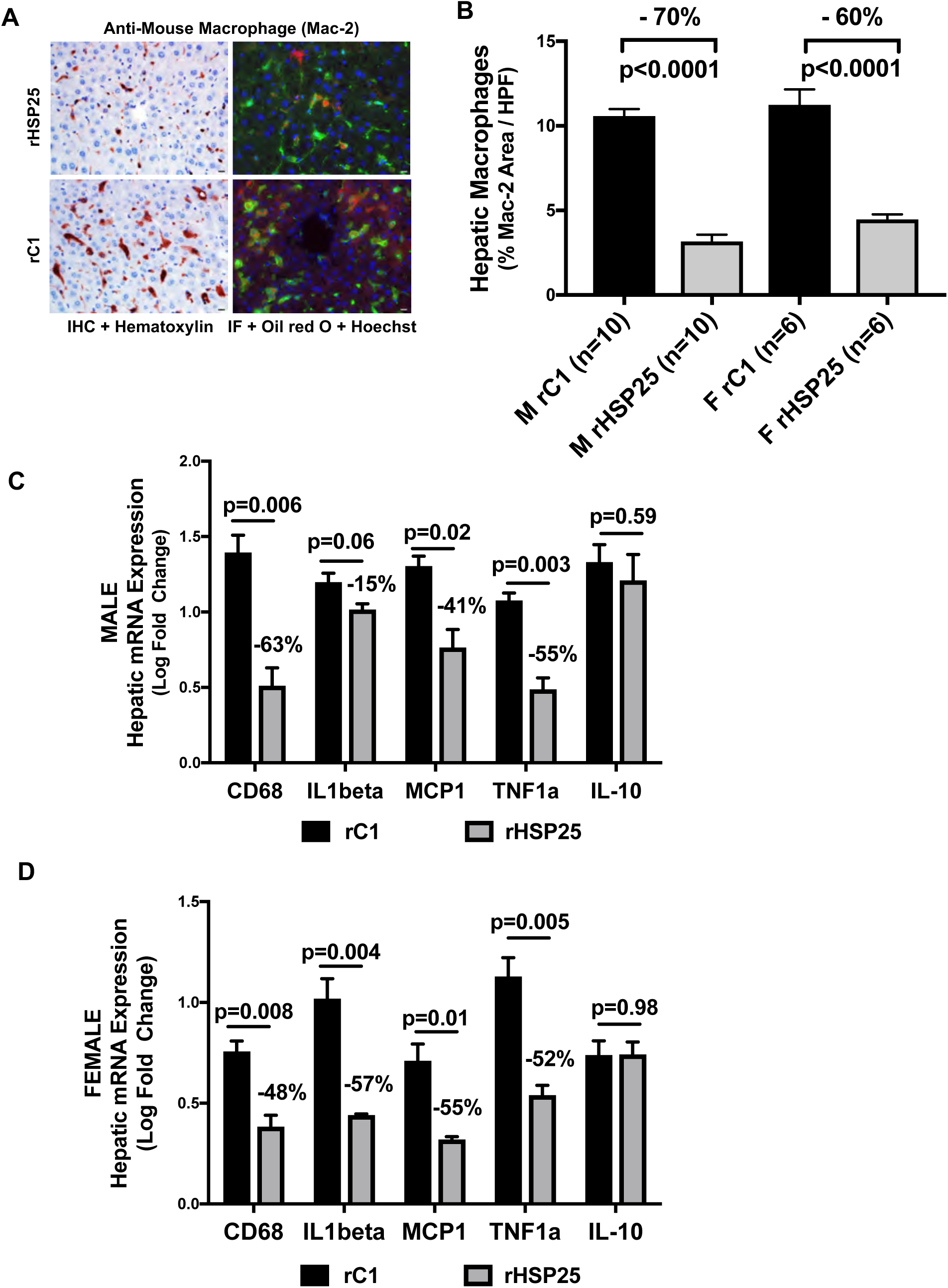
rHSP25 Vaccination reduces Hepatic Inflammation A) and B) Vaccination with rHSP25 *vs.* rC1 resulted in a marked reduction in hepatic M*Φ* content. Mac-2 immunolabeling represented by brown color, Mac-2 immunofluorescence (IF) yields a green color, the lipid deposits are red because of the oil red O stain and the nuclei have a blue Hoechst stain. Scale bar = 10 µm. C) and D) Hepatic tissue from *ApoE^-/-^* mice vaccinated with rHSP25 (*vs*. rC1) showed reduced expression of inflammatory markers / cytokines: CD68, IL-1*β*, MCP1 and TNF1*α*, with no change in IL-10 expression in this sex-disaggregated analysis.

### rHSP25 Immunotherapy Reduces Plasma Cholesterol and PCSK9 Levels and Markedly Increases LDLR Expression

Total plasma cholesterol levels were decidedly reduced in mice treated with rHSP25 relative to rC1: −59% in male (1,355 ± 113 *vs*. 552 ± 48 mg/dl, n=10 per group; p<0.0001) and −57% in female mice (1,086 ± 114 *vs.* 467 ± 36 mg/dl, n=8 per group; p=0.0001) (**Fig. 4A**). Of note, mice maintained on the HFD for an additional 5 weeks after the last rHSP25 injection continued to show a persistent reduction (−54%) in plasma cholesterol levels compared to rC1-treated control mice (**Supplemental Fig. 2C**).

**Fig. 4.**
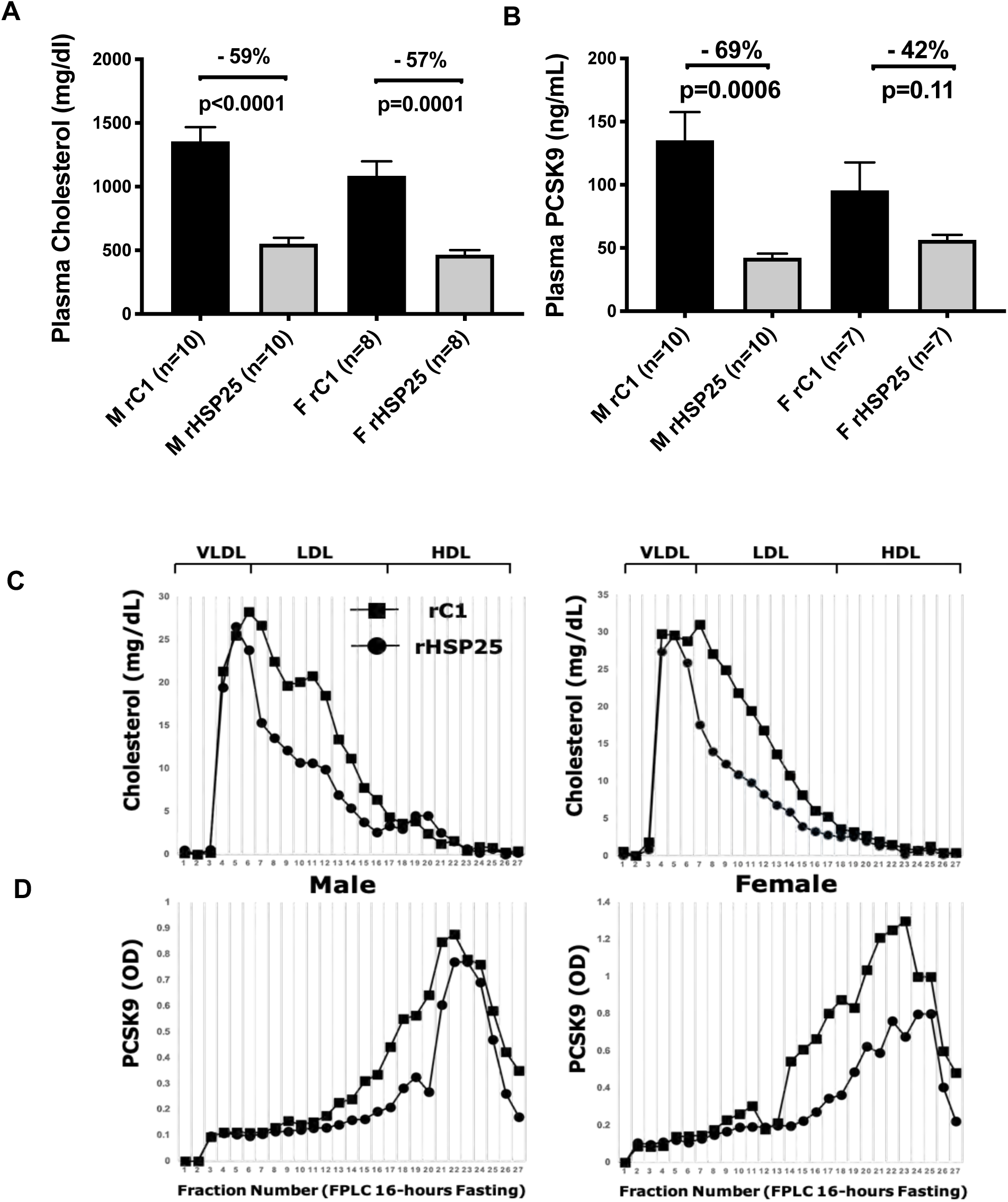
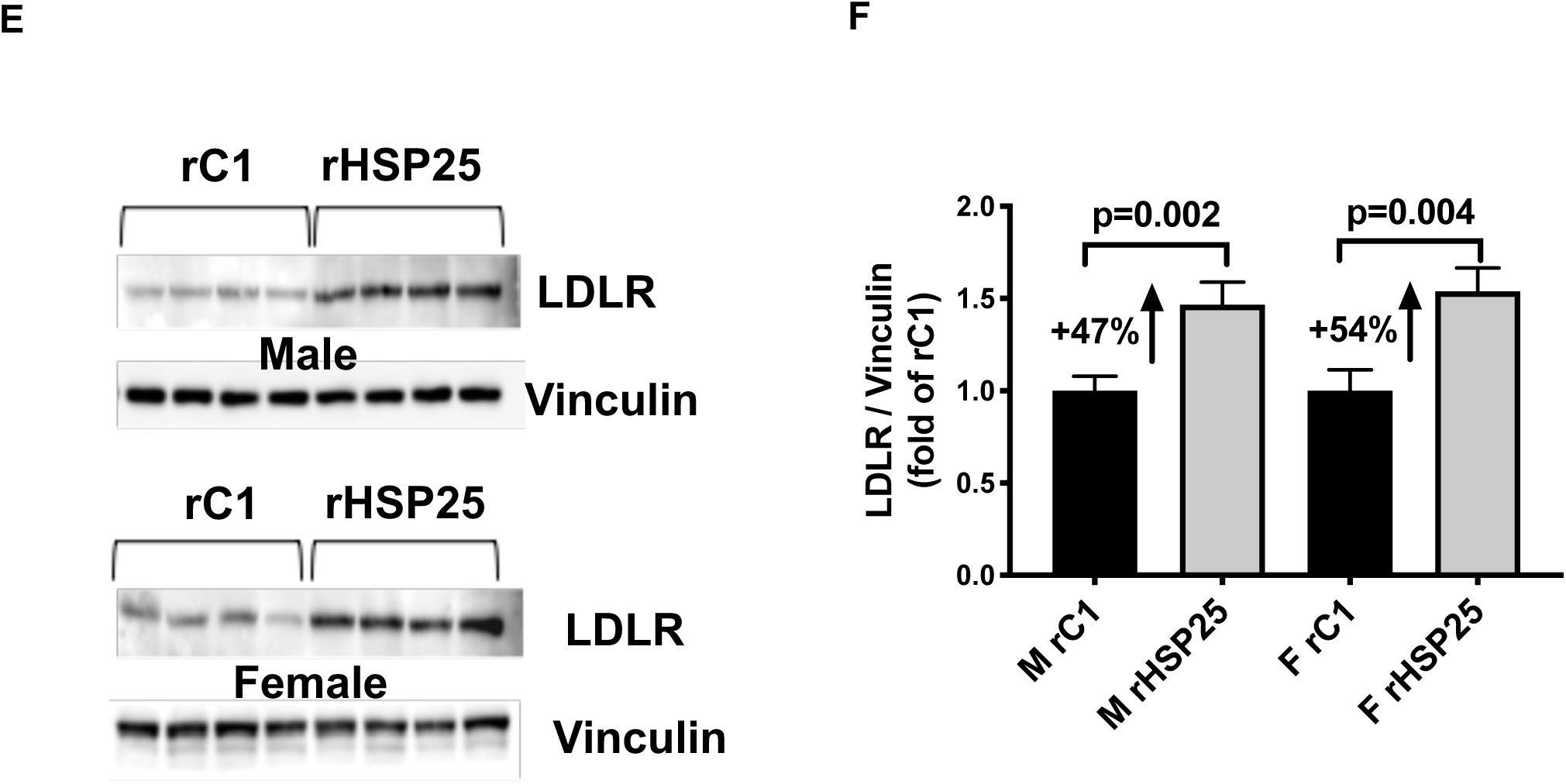
rHSP25 vaccination of *ApoE^-/-^* mice lowers plasma cholesterol and PCSK9 levels. A) At the completion of the study there was a marked decrease in plasma cholesterol levels in mice treated with rHSP25 *vs.* rC1. B) rHSP25 treatment reduces plasma PCSK9 levels in males but the reduction in females was not significant. C) and D) Cholesterol (top) and PCSK9 (bottom) measured by FPLC in pooled plasma fractions from male and female mice (rC1: n=3/group; rHSP25: n=3/group). Lipid sub-fractions based on size are indicated above the cholesterol graphs. In addition to noting reductions in the LDL-C subfraction, rHSP25 treatment reduced PCSK9 moieties for both the male and female mice. E) - F) Compared to rC1 vaccination with rHSP25 resulted in higher levels of hepatic LDLR protein expression as assessed by Western blotting in both male and female *ApoE^-/-^* mice. Each band represents protein expression for an individual mouse, and each bar graph denotes the average LDLR band intensity (normalized to the housekeeping protein vinculin) for each sex-specific treatment group.

To investigate potential mechanisms by which rHSP25 vaccination reduces atherosclerosis and plasma cholesterol levels, we measured plasma PCSK9 levels. Treatment with rHSP25 produced reductions in measured plasma PCSK9 levels in male mice (−69%; 135 ± 22 *vs*. 42 ± 11, n=10/group; p=0.0006), with a trend toward significance in female mice (−42%; 96 ± 22 *vs*. 56 ± 11, n=7/group; p=0.11) (**Fig. 4B**). When subjected to size exclusion fast protein liquid chromatography (FPLC) separation, the plasma from the rHSP25 vaccinated mice show a clear reduction in the LDL-C fraction (and in males there was a slight increase in HDL) compared to rC1 vaccinated mice (**Fig. 4C**). PCSK9 levels in the FPLC elution fractions were lower with rHSP25 vaccination in both male and female mice (**Fig. 4D**). Total cholesterol levels were directly related to PCSK9 levels (r^2^ = 0.380; p=0.0008).

Interestingly, hepatic PCSK9 mRNA and protein expression was similar in the rC1 compared to rHSP25 treated mice (**Fig. S3A – S3C**), thereby raising the possibility that the reduction in plasma PCSK9 levels may be mediated via non-transcriptional / non-translational mechanisms. Moreover, protein expression for either Sterol Regulatory Element-Binding Protein 2 (SREBP2) or Hepatocyte Nuclear Factor 1-alpha (HNF1*α*), two key transcriptional regulators of PCSK9 (**Fig. S3D, S3E**)^18, 19^ were unchanged with vaccination. Finally, there was a definite increase in hepatic LDLR protein expression in the rHSP25-*vs*. rC1-treated mice (males: 47%; p=0.002; females: 54%; p=0.004; **Fig. 3E, 3F**). Importantly, hepatic lipid content appeared diminished with rHSP25 compared to rC1 vaccination (**Fig. S3G**). Moreover, as reflected by hepatic histology, plasma aspartate aminotransferase or glucose levels, there was no evidence of off-target toxicity with rHSP25 treatment (**Table S2**).

### HSP27 Regulation of PCSK9 and LDLR Expression in HepG2 Cells

As rHSP25 vaccination resulted in striking reductions in cholesterol, with modest changes in PCSK9 levels but a robust increase in LDLR expression, we examined the potential role of the HSP27 IC in controlling the expression of key cholesterol regulatory proteins. Using Stable Isotope Labeling of Amino Acids in Cell culture (SILAC) followed by Mass Spectrometry (MS) the proteomic profile on HepG2 cells was quantified after treatment with the HSP27 IC. First, to form the IC we had to generate and validate a polyclonal anti-HSP27 IgG antibody (PAb).

Although the PAb was generated in rabbits, it recognized the same HSP27 epitopes as human AAbs from either normal subjects or CAD patients (**Supplemental Fig. 4A-4D**). For the SILAC experiment, control HepG2 cells were cultured in ‘light’ media, while cells treated with [rHSP27 + PAb] were cultured in media containing ‘heavy’ isotope-labelled amino acids. The ratio of ‘heavy’ to ‘light’ proteins synthesized with these treatments was normalized to that of GAPDH, and therefore reflects the relative changes in protein expression due to treatment with the HSP27 IC. Treating HepG2 cells with 1 µg/ml rHSP27 combined with 5 µg/ml PAb (∼1:1 molar ratio) resulted in a 33% decrease in PCSK9 (p<0.0001) and an 44% increase in LDLR (p=0.0009) levels, with no effect on other known modulators of cholesterol metabolism, such as HMG Co-A reductase (HMGCR), SREBP2, or the Inducible Degrader of the LDL receptor (IDOL) (**Fig. 5A**).

**Fig. 5.**
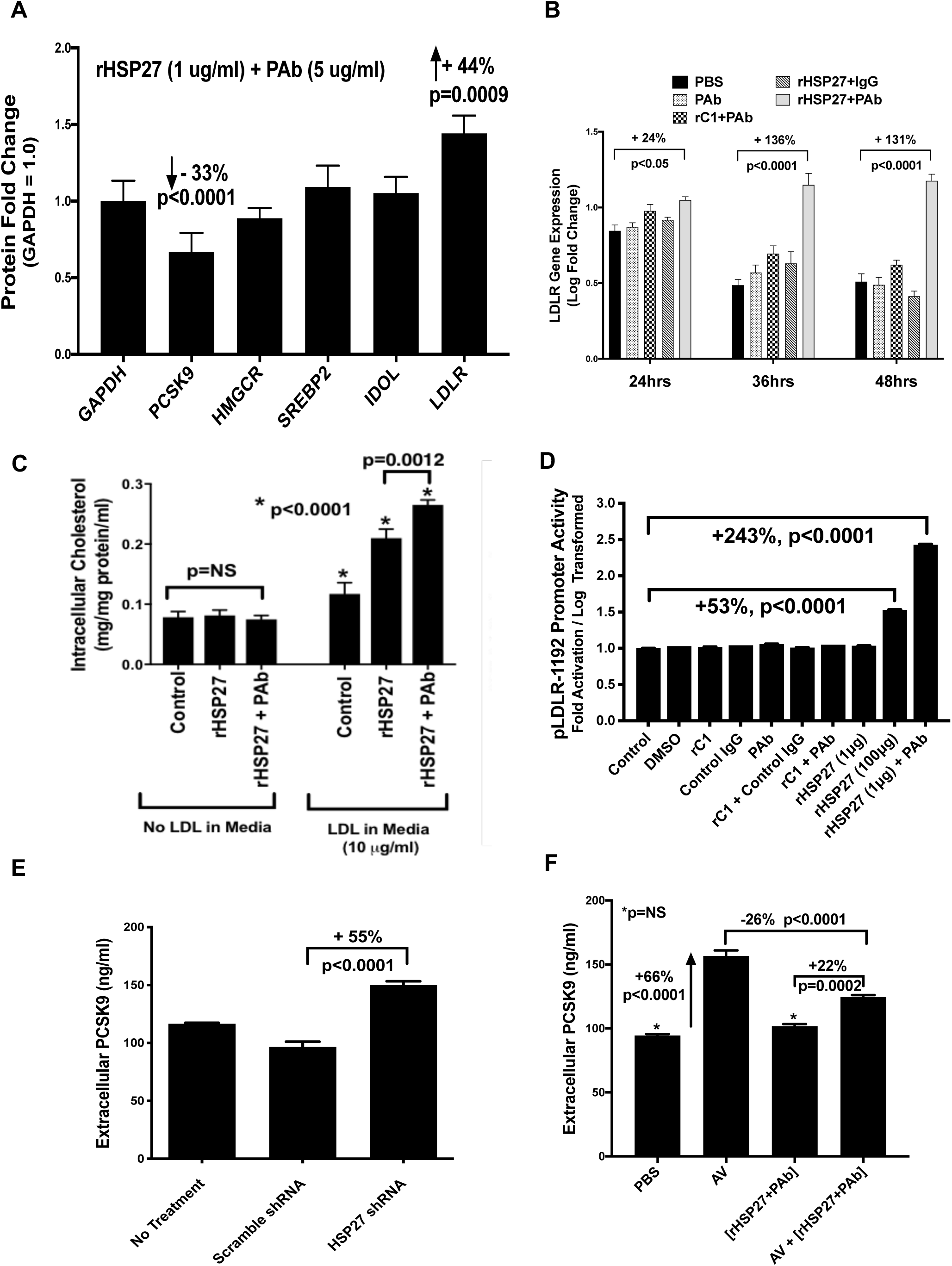
HSP27 plus anti-HSP27 polyclonal antibody (PAb) modulate PCSK9 and LDLR in HepG2 cells. A) SILAC experiment performed after 18 hrs treatment with [rHSP27 (1 µg/mL) + PAb (5 µg/mL)]. The relative abundance of proteins involved in cholesterol metabolism was expressed as the ratio of their abundance with treatment *vs.* control conditions. Each ratio was then normalized to the ratio for the housekeeping protein GAPDH. Compared to control, the combination of [rHSP27 + PAb] decreased PCSK9 and increased LDLR expression. B) rHSP27 plus AAb increase LDLR gene expression. Time course of changes in LDLR gene expression post-treatment with [rHSP27 (1 µg/mL) plus PAb (5 µg/mL)] or control treatments [PBS, PAb alone (5 µg/mL), rC1 (1 µg/mL) plus PAb (5 µg/mL), and rHSP27 (1 µg/mL) plus non-specific polyclonal IgG (5 µg/mL)]. Gene expression was assessed by qPCR and presented as log fold change. Relative to control treatments [rHSP27 + PAb] produced profound and sustained increases in LDLR expression. C) The increase in LDLR expression occurs without a drop in intracellular cholesterol levels. In fact, there is an increase in intracellular LDL-C concentration when HepG2 cells are incubated with LDL-C (10μg/ml) in the media. D) HepG2 cells transfected with a full length LDLR promoter construct (pLDLR-1192, extending from −989 to +203 relative to the transcription start site of human LDLR gene) and employing firefly luciferase to demonstrate promoter activation. Various controls and treatments are outlined, with the results expressed as log transformed activation. Compared to control LDLR promoter activity was strikingly increased with [rHSP27 + PAb] treatment. E) Extracellular PCSK9 protein levels are reduced in HepG2 cells treated with small hair pin RNA (shRNA) sequences designed to knockdown the expression of HSP27. No treatment and treatment with a scrambled shRNA represented control conditions. Cell supernatant levels of PCSK9 protein increased when HSP27 expression was knocked down *vs.* scrambled shRNA sequence (quantification by Western blot with band densitometry using *β*-actin as the housekeeping protein). F) Overnight treatment of HepG2 cells with AV (10μM) increased PCSK9 levels in the cell supernatant, while [rHSP27 (1 µg/mL) plus PAb (5 µg/mL)] had no effect on its own, but did attenuate the AV-increase.

Next, the potential role of the HSP27 IC in regulating expression of PCSK9 and its transcriptional factor HNF1*α* was examined in HepG2 cells treated with: [rHSP27 (1 µg/mL) plus PAb (5 µg/mL)], PBS, PAb alone (5 µg/mL), [rC1 (1 µg/mL) + PAb (5 µg/mL)], [rHSP27 (1 µg/mL) + non-specific rabbit IgG (5 µg/mL)], with the results reported as log-fold change. Relative to PBS, treatment with PAb alone, [rC1 + PAb], and [rHSP27 + control IgG] failed to show a significant and persistent alteration in PCSK9 or HNF1α expression (**Supplemental Fig. 5A - 5B**). For example, [rHSP27 + PAb] treatment resulted in a transient 27% reduction in PCSK9 gene expression compared to PBS treatment, however this only occurred at 6 hrs (p<0.001). There were slightly larger and persistent reductions in HNF1α gene expression with [rHSP27 + PAb] treatment – but again these were transient (e.g., −36% at 6 hours, −35% at 12 hours, p<0.0001 for both).

In contrast, LDLR expression in HepG2 cells was intensely upregulated with [rHSP27 + PAb] treatment and for a protracted interval: at 24 hours: +24% (p<0.05), 36 hours: +136% (p<0.0001), and 48 hours: +131% (p<0.0001) (**Fig. 5B**). Normally, LDLR expression is regulated by intracellular cholesterol concentrations (i.e., via SREBP2); however, both *in vivo* (**Supplemental Fig. 3D**) and *in vitro* (i.e., SILAC experiment; **Fig. 5A**) SREBP2 protein expression remained unchanged with rHSP25 vaccination and [rHSP27 + PAb] treatments.

Indeed, this increase in HepG2 LDLR expression occurred without a reduction in intracellular cholesterol concentration. Treatment with [rHSP27 + PAb] produced an increase in LDL-C uptake and intra-cellular cholesterol concentration (**Fig. 5C**). Nonetheless, *in vivo* there was no evidence of excessive intra-hepatic accumulation of lipid (**Fig. 3A, Supplemental Fig. 3F**).

LDLR promoter activity was assessed using the plasmid pLDLR-1192, containing the full length promoter region from −989 to +203 relative to the transcription start site of human LDLR gene, as well as the plasmid pLDLR-234, containing the LDLR core promoter sequence with only the Sterol Response Element-1 (SRE-1) and Sp1 regulatory element sites from −142 to +35^20^. Compared to control treatment, the full length LDLR promoter showed a 243% increase in activity after treatment with low dose [rHSP27 (1μg) + PAb] compared to a 53% increase with high dose (100μg) rHSP27 alone (**Fig. 5D**). The results with the LDLR core promoter construct were less (e.g., +46% with [rHSP27 (1μg/ml) + PAb] and +23% with 100μg of rHSP27 alone), thereby suggesting that there may be a novel regulatory mechanism of LDLR transcription that is upstream and absent in the core promoter construct (**Supplemental Fig. 5C**).

Finally, to further assess the role of HSP27 on PCSK9 expression, HepG2 hepatocytes were transfected with a previously described short hairpin RNA (shRNA) targeting HSP27 (**Supplemental Fig. 5D**)^21^. Silencing HSP27 resulted in 55% higher levels of PCSK9 in the culture media compared to a scrambled control sequence (p<0.0001; **Fig. 5E**). To determine if the HSP27 IC could attenuate a statin-induced rise in PCSK9, HepG2 cells were treated with atorvastatin (AV; 10μM), resulting in a 66% increase in PCSK9 mRNA *vs.* PBS control (**Fig. 5F**). On its own [rHSP27 + PAb] was similar to PBS control. Adding [rHSP27 + PAb] dampened the AV-induced increase in PCSK9 mRNA by 26% (p<0.0001).

### LDLR Expression is mediated via the NF-κB Pathway

In the past, we repeatedly demonstrated in M*Φ* that HSP27 activates the NF-*κ*B pathway to alter the transcription of several key genes that are either pro- or anti-atherogenic^21–24^. We showed in M*Φ* that activation of NF-*κ*B by HSP27 required TLR4 and could be blocked with various inhibitors of the NF-*κ*B intracellular pathway (e.g., CLI-095 an inhibitor of TLR4; IRAK1/4 an inhibitor of the TLR4 pathway downstream secondary messenger IRAK; MG132 to block the proteasomal degradation of I*κ*-*β*)^21^. However, we did not explore if NF-*κ*B activation by the HSP27 IC is integral to LDLR expression, and now present the following data. First, we illustrated the activation of the NF-*κ*B pathway in hepatocytes, observing a strong immunofluorescent nuclear localization signal for the NF-*κ*B p65 subunit in human hepatic cells treated with [rHSP27 + PAb] and much less so after treatment with rHSP27 (alone) or the PAb (alone) (**Fig. 6A**). Second, to explore the potential role of NF-*κ*B in the upregulation of LDLR expression by HSP27, HepG cells were pre-incubated with (or without) the NF-*κ*B pathway inhibitor, BAY 11-7082 for 30-60 minutes before being treated for up to 24 hours with [rHSP27 + PAb]. Compared to control, LDLR mRNA expression was increased by [rHSP27 + PAb] (66%; p<0.0001), and reduced by the BAY 11-7082 alone (35%, p=0.0008; **Fig. 6B**). Moreover, BAY 11-7082 inhibited the augmentation effect of [rHSP27 + PAb] on LDLR mRNA expression, keeping them at levels similar to control treatment. Third, relative to controls, the HSP27 IC increased LDLR protein expression by 70% at 16 hrs and 40% at 24 hrs (p=0.0008 and p=0.016; respectively) – escalations that were essentially nullified in the presence of BAY 11-7082 alone or combined with [rHSP27 + PAb] (**Fig. 6C – 6D**).

**Fig 6.**
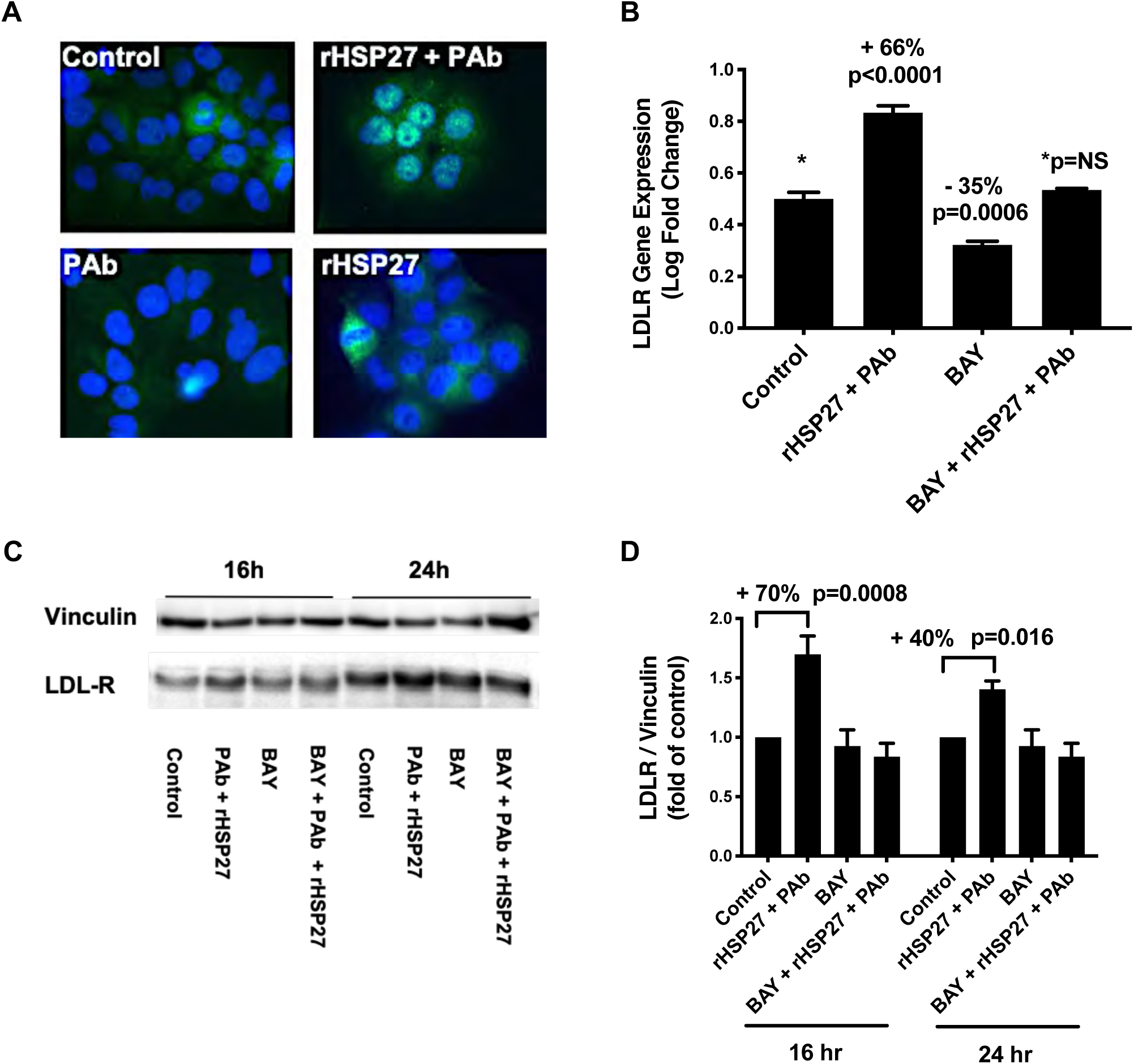
HSP27 Upregulates LDLR Expression via NF-*κ*B Activation. A) Human hepatocytes treated with rHSP27 (1μg/ml) with or without PAb (5 μg/ml) for 45 minutes before being labeled with a fluorescent rabbit anti-NF-κB antibody and a Hoechst nuclear label. A fluorescent nuclear localization signal was most abundant when cells were treated with [rHSP27 + PAb] (original photomicrograph: *×*400 magnification). B) In HepG2 cells LDLR gene expression increased with [rHSP27 + PAb] treatment. The addition of NF-*κ*B blocker BAY11-7082 reduced basal LDLR expression by 35% *vs.* Control, and annihilated the increase induced by [rHSP27+PAb]. C) – D) Treatment of HepG2 cells for 16 and 24 hrs with [rHSP27 + PAb] increased LDLR protein expression – effects that were blocked by co-treatment with BAY 11-7082.

## Discussion

The current clinical studies demonstrate that anti-HSP27 IgG blood levels are elevated in healthy subjects compared to patients with CVD and confirm our previous report HSP27 blood levels are also higher in health^11^. To explore the role of these AAbs in atherosclerosis, *ApoE^-/-^* mice underwent rHSP25 vaccination resulting in the attenuation of the early inflammatory stages of atherogenesis, with less intra-plaque cholesterol accumulation, as well as reduced plaque and hepatic inflammation. These findings are different from our previous studies in *ApoE^-/-^* mice injected twice daily with rHSP27 because: i) we used an adjuvant to purposely generate anti-HSP25 antibodies, ii) the treatment schedule was reduced to once weekly injections, and iii) the reductions in plasma cholesterol levels were more profound (∼60% *vs*. 42%) and persistent (**Supplemental Fig. 2C**)^11^.

Anti-HSP27 antibodies are natural antibodies produced in the absence of infection or immunization, differ from adaptive antibodies in repertoire and function, and are thought to decline in abundance with age^25^. Natural antibodies can consist of various species (e.g., IgM, IgA and IgG), and may provide primary protection against infection during the gap prior to germinal center formation and adaptive antibody production^26^. While natural antibodies may also be auto-reactive and assist with homeostatic housekeeping tasks, removing cellular debris and noxious species such as oxidized LDL-C^27–29^, they do not necessarily destroy their targets^30^.

Indeed, not all natural antibodies are antagonistic^31–33^ and antibody titers of any nature may represent some form of health advantage. For example, in the large Anglo-Scandinavian Cardiac Outcome Trial (ASCOT), elevated levels of IgG (regardless of the recognized antigen) are associated with a reduced risk of coronary heart disease^32^. While we also observed increased IgM levels, unlike IgG AAb they did not independently associate with either health or CVD. Currently, we are exploring how these IgG AAbs facilitate HSP27 signaling, interacting with TLR4 to bind at the cell surface and activate NF-*κ*B.

To study the mechanisms by which the HSP27 IC reduce plasma cholesterol levels, we examined the expression of two key regulators of LDL-C levels, PCSK9 and LDLR. rHSP25 vaccination reduced plasma PCSK9 levels (**Fig. 4B, 4D**) but prompted little change in hepatic PCSK9 mRNA and protein expression (**Supplemental Fig. 3A-3C**). However, *in vitro* treatment of human hepatocytes with [rHSP27 + PAb] reduced PCSK9 protein expression by 33% (SILAC experiment; **Fig. 5A**) but without impressive changes in mRNA expression (**Supplemental Fig. 5A**). Interestingly, rHSP25 vaccination or the HSP27 IC *in vitro* had only minor effects on the expression of SREBP2 and HNF1*α*, the two transcriptional regulators of PCSK9 (**Supplemental Fig. 3D, 3E, 5B**).

In contrast, there was a marked increases in hepatic LDLR protein expression with rHSP25 vaccination [+47% in male and +54% in female mice; **Fig. 4E, 4F**), with complementary changes *in vitro* (+44% increase in LDLR protein in the SILAC experiment, +136% upregulation of LDLR mRNA levels, +243% increase in LDLR promoter activity with [rHSP27 + PAb] treatment; **Fig. 5A, 5B, 5D**). Interestingly, we noted no changes in total plasma cholesterol levels in *LDLR^-/-^* mice vaccinated with rHSP25 – thereby supporting the concept that rHSP25 immunotherapy requires LDLR to effect atheroprotection (**Supplemental Fig. 2A**). LDLR is the primary determinant of not only LDL-C^3^ but also PCSK9 levels – as the LDLR is the principal clearance mechanism for plasma PCSK9^34^. Considering that the LDLR recycles every 10 minutes, internalizing hundreds of LDL-C particles during its ∼20 hrs lifespan^35^, small increases in LDLR expression translate into large decreases in LDL-C as well as PCSK9 removal from the plasma. Finally, knocking down HSP27 expression increases extracellular PCSK9, while adding [rHPS27 + PAb] to HepG2 cells attenuates statin-induced increases in extracellular PCSK9 levels (**Fig. 5E, 5F**).

In general, hepatocyte LDLR levels are thought to be regulated at the transcriptional level by a SRE in the LDLR promoter. In response to low intracellular cholesterol levels (e.g., with statin therapy), SREBP2 activity increases and binds to this SRE. However, other mechanism may also alter LDLR expression (e.g.,ubiquitin-induced proteasomal degradation of HNF1*α* by berberine, or a PPAR-response element in the LDLR promoter)^20, 36^. *In vitro,* we note that intracellular cholesterol levels actually rise with rHSP27 IC treatment (with no corresponding hepatic steatosis or toxicity *in vivo*) (**Fig. 3A, 5C; Supplemental Fig. 3F, Supplemental Table 2**) and do not alter SREBP2 levels (**Supplemental Fig. 3D**). Hence, unlike statins, which result in increased expression of both LDLR and PCSK9, the effect of [rHSP27 + PAb] appears to be divergent, reducing PCSK9 levels (e.g., by a direct transcriptional effect and/or indirectly via LDLR clearance of PCSK9) yet promoting a marked increases in LDLR promoter activity and mRNA/protein expression.

How HSP27 upregulates LDLR expression is also becoming better understood. Previously we demonstrated that extracellular HSP27 (alone) can activate intracellular signaling cascades such as NF-*κ*B via TLR4 in M*Φ*^21–23^. While NF-κB signaling is associated with pro-inflammatory responses in atherosclerosis^37, 38^, it also regulates anti-inflammatory processes^39^ and deletions of components of this pathway aggravate inflammation^40–42^. Hence, NF-*κ*B signaling is likely very much influenced by context, with the tip of the inflammation balance determined by cell type and tissue distribution. Interestingly, we now show for the first time that [rHSP27 + PAb] activates NF-*κ*B in hepatocytes (**Fig. 5A**) and that this is a required step for the upregulated expression of LDLR mRNA (66%) and protein (70%) (**Fig. 6B-6D**). Collectively, these data point to a previously unrecognized LDLR regulation pathway that may be druggable.

Our study has limitations. For example, because of the prevalence of statin use in our clinical cohort from Alabama (CVD: 96% and CON 17%; **Table S1**) we are unable to determine a meaningful relationship between the LDL-C (or PCSK9) and HSP27 or AAbs. As well, we have separately quantified human levels of HSP27 and AAbs (**Fig. 1A, 1B**) but not levels of the HSP27 IC. However, in pooled human blood samples we document the joint presence of IgG immunoglobulins and HSP27 (**Supplemental Fig. 6A**) but, as this involved Western blotting we cannot provide a full quantitative comparison of the IC abundance in CON *vs*. CVD subjects. Furthermore, using the Duolink Proximity Ligation Assay, we graphically illustrate the interaction between HSP27 and PAb in hepatocytes (**Supplemental Fig. 6B**).

In summary, HSP27 immunotherapy is a promising new strategy to reduce atherogenesis by lowering cholesterol levels through the upregulation of LDLR expression that promotes enhanced clearance of LDL-C. Given the rapid recycling of the LDLR, we postulate that this increase in LDLR synthesis augments PCSK9 intracellular disposal. In addition, the observation that [HSP27 + PAb] increases LDLR expression in an NF-*κ*B dependent manner that is independent of intracellular cholesterol levels is novel and uncovers an attractive opportunity for the development of a new therapeutic avenue for managing cholesterol disorders. Going forward, we know that low HSP27 and AAb levels are associated with CVD, and are focusing our efforts on developing therapies to boost AAb levels in order to potentiate existing (low) levels of HSP27. Having mapped the HSP27 epitopes (**Supplemental Fig. 4A –4C**) we are striving to optimize the efficacy of HSP27 vaccination for a more prolonged therapeutic benefit. With other HSP27 immunological approaches also under consideration, (e.g., the direct administration of anti-HSP27 monoclonal antibodies for passive immunization), we anticipate important advances in treating both dyslipidemia and inflammation; hence targeting atherogenesis.

## Acknowledgments

We are indebted to the Libin Cardiovascular Institute of Alberta and its community partners for supporting the <u>O’Brien Vascular Biology Laboratory</u>, as well as the staff in the University of Calgary Cumming School of Medicine Health Sciences Animal Resource Centre.

## Clinical Study of Serum HSP27 and Anti-HSP27 Antibody Levels

As part of a National Institute of Health sponsored project (#5P20MD002314-08; Sub-Project ID: 6374) designed to identify cardiovascular biomarkers for ischemic heart disease in a population that is medically underserved, subjects were enrolled in a 5-year follow-up study that involved the collection of blood samples at various University of South Alabama-affiliated medical clinics. The study was approved by the University’s Institutional Review Board and complied with the Declaration of Helsinki. In total, 267 cardiovascular disease (CVD) patients aged 45-80 years of either sex were recruited from the Heart Center, University of South Alabama Health System, Mobile, AL as were 100 healthy volunteers (CON; either patients or University of South Alabama employees). The demographics of the clinic population was estimated to include approximately 40% medically underserved individuals and 27% African-Americans.

Using a nested case control strategy, CVD patients were identified and compared to CON with patient characteristics blinded to investigators for all assays. The enrollment criteria for the CVD patients included any of the following: typical symptoms of angina pectoris lasting >20 minutes at rest, ECG changes during angina symptoms, hypertension, a diagnosis of an acute coronary syndrome or a history of coronary revascularization. Patients were excluded if they suffered from renal impairment, autoimmune disease, or rheumatic heart disease – all factors known to alter levels of inflammatory biomarkers, including heat shock proteins. The CON cohort lacked any of the inclusion or exclusion criteria. For the current study, 80 CVD and 58 CON subjects were blindly and randomly selected. Relevant clinical characteristics are described in **Supplemental Table 1**.

## Statistical analyses for Clinical Study

For the clinical data, continuous variables are reported as mean ± SD. Categorical variables are reported as number (%) and were compared using χ2 or Fisher’s exact tests. Chi-square tests were performed on nominal data sets: male sex, smoker, diabetes mellitus (defined by use of hypoglycemic agents), hypertension (defined by of conventional anti-hypertensive therapy), statin use (defined by use of any HMG Co-A reductase medication), African American and Caucasian. All comparisons used two-tailed alpha levels of 0.05. In cases of multiple comparisons, one-way analysis of variance (ANOVA) and Tukey’s method were used to correct for the family-wise error rate. The association of health *vs.* CVD status with blood concentrations of HSP27 and AAb was examined using multivariable linear regression, adjusting for age, sex, ethnicity, body mass index (BMI), diabetes mellitus, and active smoking status. Adjusted differential levels in HSP27 and AAb are reported with 95% confidence intervals (CI). All analyses were performed in a blinded fashion using GraphPad Prism 8 (GraphPad Software, La Jolla, CA), except for regression models, which were fit using SAS v9.4 (SAS Institute Inc., Cary, NC).

## Recombinant protein preparation

Plasmids encoding HIS-tagged full-length HSP27, the mouse ortholog HSP25, or the C-terminal, biologically inactive (rC1) fragment of HSP27 (AA93-205) were constructed using a pET-21a vector, with the plasmids transformed into an *Escherichia coli* expression strain (DE3) as described previously.^1^ Recombinant proteins were purified with a Ni-NTA resin and Q-Sepharose^TM^ (GE Healthcare). Endotoxin was removed by Pierce High-Capacity Endotoxin Removal Resin (ThermoFisher Scientific, Waltham, MA, USA). The purity of the final recombinant proteins was determined to be more than 99% by SDS-PAGE with an endotoxin concentration lower than 2 units/mg protein measured by Limulus Amebocyte Lysate PYROGENT^TM^ 125 Plus (Lonza).

## Measurement of HSP27 and antibodies to HSP27 (AAbs) Levels in Human Sera

Blood HSP27 levels were measured in plasma samples using an enzyme-linked immunosorbent assay (ELISA; Human HSP27 DuoSet ELISA, DY1580, R&D systems) previously reported by our group.^2, 3^ The assay detection range was 31.3 to 2,000 pg/ml. The average intra-assay coefficient of variation (CV) was <6.3% and the inter-assay CV was 10.3%. The absorbance was measured at 450 nm with a microplate reader (Synergy Mx, BioTek, Winooski, VT, USA).

IgG and IgM AAb levels were measured using an ELISA developed in our laboratory. Briefly, NUNC maxisorp plates (ThermoFisher) were coated with rHSP27 at a concentration of 500 ng/well in carbonate-bicarbonate buffer overnight (ON) at room temperature (RT). The wells were blocked with 1% BSA/PBST, washed in phosphate buffered saline tween (PBST), and incubated with plasma at a final dilution of 1:2,000 in 1% BSA for two hrs followed by 3 more washes in PBST. A horse radish peroxidase (HRP) labeled anti-human IgG (H&L) antibody #109-035-003, Jackson Immunoresearch, West Grove, PA) was used as a detection antibody at a dilution of 1:5,000 and incubated for one hr at RT. Finally, substrate solution (3,3’,3.5’-Tetramethylbenzidine Liquid Substrate, TMB; Millipore Sigma) was added to each well and incubated for 10 mins avoiding direct light. The reaction was stopped by 2N H_2_SO4 and the optical density quantified at 450 nm using Synergy Mx plate reader (BioTek). The detection of IgM antibodies to HSP27 was performed in a manner similar to that used for the detection of the anti-human IgG described above, with the exception that a goat anti-human IgM antibody, Fc5μ, HRP conjugate (AP114P, Millipore, Sigma) was substituted in place of the anti-human IgG antibody. To establish an internal standard for the measurement of HSP27 auto-antibodies, plasma from a healthy control subject was diluted 500 times in 1% BSA/PBST and arbitrarily defined as 50 absorption units (a.u.). For the murine model described below, AAb blood levels were measured similar to the human assay, except rHSP25 replaced rHSP27 for the coating the NUNC maxisorp plates.

To simultaneously detect the presence of HSP27 and AAbs blood samples from healthy subjects or CVD patients were (separately) pooled (i.e., samples from 6 healthy subjects *vs.* 6 CVD patients, with three biological replicates per group). Blood samples were prepared using an microvesicle isolation protocol that we previously published^4^. Briefly, we used ultracentrifugation cycles of 2,000*×*g, 10,000*×*g, 100,000*×*g to obtain a pellet that contained microvesicles. We then resolved the resultant pellet using SDS-PAGE followed by Western Blotting (**Fig. S1D**). From this, we could detect HSP27 and (separately) human IgG in the pellet fraction of the plasma which contains microvesicles.

## Murine Model for the Study of Atherosclerosis

Male and Female ApoE^-/-^ mice (C57BL/6 genetic background, Jackson Laboratory, Bar Harbor, ME, USA; stock 002052: B6.129P2-*Apoe^tm1Unc^*/J) were fed a normal chow diet (9% fat) until 8 wks of age, followed by a high fat diet (HFD, 15.8% fat, 1.25% cholesterol, Envigo-Teklad diet #TD94059) for 6 wks until euthanasia. After a 2 wk run-in period with HFD, mice were randomized to vaccination with either rHSP25 (M: n=10, F: n=8) or rC1 (M: n=10, F: n=8), or PBS subcutaneous injections weekly for 4 wks then euthanized. More precisely, vaccination was performed by mixing rHSP25 (100 µg, 75 µL) or rC1 (100 µg, 75 µL) with an aluminum hydroxide gel adjuvant (Alhydrogel® 2%, Al, 25 µL; Invivogen vac-alu-250, San Diego, CA). The blood samples were collected (EDTA tube) weekly from the saphenous vein and stored at −80°C for future use. At the end of the study blood samples were obtained via the right heart ventricle after 16 hrs of ON fasting. This was followed by whole body perfusion with PBS. The left liver lobe was then flash frozen in liquid nitrogen and stored at −80°C for gene and protein expression assays. Thereafter, mice were systemically perfused with 10% NBF via the heart left ventricle, and the heart, aorta and liver tissues were removed and immersed in 10% NBF ON. In addition, we performed this same experiment in: i) LDLR^-/-^ mice (C57BL/6 background, Jackson Laboratory, Bar Harbor, ME), and ii) apoE^-/-^ mice with the *Hspb1* gene knocked out (*Apoe^-/-^ HSP25^-/-^* mice, generated by J.L.E. Dean, Oxford, UK and previously described.^5^ Levels of AAbs in murine plasma were measured in a manner similar to that used for the clinical samples, but employed a goat anti-mouse IgG (H&L) antibody (catalogue #115-035-062) that has minimal cross reactivity against human epitopes. All experimental murine procedures were approved by the Animal Care Committee of the University of Calgary.

## Evaluation of *en face* Atherosclerotic Lesion Area and Hepatic Lipid Content

The aorta was dissected from the ascending to the thoracic segments ending at the 4^th^ bronchial branches and the adventitia was removed. With the primary incision following the lesser curvature of the arch, the aorta was then opened longitudinally, and a second incision was made along the greater curvature of the arch down to the level of the left subclavian artery. As described previously, the aortic and hepatic lipid content was evaluated on oil red O stained slides^6^ and photographed using an Olympus SZ-61 stereo microscope with an Olympus UC30 camera (aortic magnification ×12, hepatic magnification *×*400). The *en face* atherosclerotic aortic lesions were analyzed using Image-Pro software (Media Cybernetics) to calculate the total and atherosclerotic lesion areas. The extent of atherosclerosis was expressed as the percentage of surface area of the entire aorta covered by lesions as described previously^3^.

## Evaluation of Aortic Sinus Lesion Area and Cleft Area

The top half of the heart containing the aortic root was embedded in paraffin or frozen in Tissue-Tek O.C.T. media. Serial 4-μm sections of the aortic sinus including the aortic valve leaflets were sectioned, beginning at the level where the aortic valve first appears, and stained with hematoxylin and eosin (H&E; Millipore Sigma, Etobicoke, ON). Atherosclerotic lesion areas were analyzed using Image-Pro software (Media Cybernetics, Silver Spring, MD). Mean lesion area for each mouse was calculated by averaging measurements from four cross-sections.

Cholesterol clefts were defined as ghost-like needle-shaped spaces that resulted from dissolution of cholesterol crystals during paraffin-embedding and tissue processing and were manually quantified using a 4*×* objective lens.^7, 8^

## Evaluation of Aortic Sinus Macrophage Content

Serial 4-μm sections of aortic sinus were blocked with 10% normal horse serum followed by rat anti-mouse macrophage primary antibody (Mac-2, Accurate Chemical and Scientific Corp.) diluted in PBS 1:500 at 4°C ON. Biotinylated rabbit anti-rat was used as secondary antibody (1:100, Vector Laboratories). Endogenous peroxidase activity was quenched with 3% H_2_O_2_.

Antibody reactivity was detected with an ABC kit (Vector Laboratories, Burlingame, California, USA) and visualized with diaminobenzidine (DAB). Sections were counterstained with hematoxylin, cleared, and mounted. Immunopositive areas in cross sections were quantified using Image-Pro software using a 4*×* objective lens.

## Liver Histology and Immunohistochemical Staining

After fixation in 10% NBF, liver tissue was embedded in paraffin, and serially sectioned at 5-μm intervals. After deparaffinization, tissue sections were rehydrated and subjected to H&E staining. Histopathological changes of liver tissue based on H&E staining were examined and photographed with an Olympus BX52 microscope. Immunohistochemical staining was performed using the following primary antibodies: rat anti-Mac-2 (1:500; CedarLane Laboratories, Burlington, ON; CL8942AP), rabbit anti-LDLR (1:200; Abcam, ab52818) and rabbit anti-PCSK9 (1:200; Abcam, ab31762). Sections of liver tissue were deparaffinized in xylene and rehydrated in graded concentrations of ethanol/water. The sections were blocked with 10% normal horse serum followed by incubation with primary antibody at 4°C ON. Peroxidase conjugated anti-rat or anti-rabbit were used as secondary antibodies (1:100, Vector Laboratories), respectively. Endogenous peroxidase activity was quenched with 3% H_2_O_2_.

Antibody reactivity was detected with an ABC kit (Vector Laboratories) and visualized with diaminobenzidine (DAB). Sections were counterstained with hematoxylin, cleared, and mounted on glass slides. Immunopositive areas in cross sections were quantified using Image-Pro software according to previously described methods ^3^. The negative control consisted of substituting IgG for the primary antibody.

## Lipid-laden Macrophage Staining in Mouse Liver Tissue

In order to interpret the lipid-laden macrophage content of liver tissue, both immunofluorescence and the lipid-soluble dye oil red O were applied on the same section, as described previously ^6^. Briefly, a 5-μm-thick tissue section was incubated in oil red O solution (Sigma Aldrich, Oakville, ON) for 1 hr at RT. Following washes in PBS, the section was pre-incubated with 10% normal horse serum in PBS for 5 minutes followed by a rat anti-mouse macrophage primary antibody (Mac-2; Accurate Chemical and Scientific Corp.) diluted in PBS 1:500 and left at 4°C ON. After repetitive rinsing with PBS, the sections were incubated with a fluorescein-green rabbit anti-rat secondary antibody (1:100, FI-4001, Vector Laboratories) for 30 minutes at RT. Following triplicate washes in PBS, tissue sections were incubated with Hoechst 33258 (Sigma Aldrich) to produce a nuclear stain. Three washes with PBS were again performed before mounting in 50% glycerol in PBS. Photomicrographs were obtained using a fluorescence microscope (Olympus BX52).

## Murine Liver Tissue Q-PCR

The expression of inflammatory M*Φ* markers and cytokines were measured in murine liver tissue: CD68, IL1*β*, MCP1, TNF1*α*, IL-10. Total RNA was isolated from a piece of the liver (100 mg) using TRIzol reagent (Thermo Fisher Scientific) and 1 µg of purified RNA was reverse transcribed to cDNA using the qScript cDNA SuperMix (Quantabio, 95048-100) mastermix.

Quantitative real-time PCR was performed at least in triplicate using PerfeCta SYBR Green Supermix ROX (Quantabio, 95055-500) on the StepOnePlus Real-Time PCR System (Thermo Fisher Scientific). *β*-Actin was used as an endogenous control to calculate fold changes of the target gene expression using the 2^−ΔΔCt^ method. In a similar manner murine hepatic PCSK9 expression was assessed but using the control gene, hypoxanthine-phosphoribosyltransferase (hHPRT1). The qPCR primers used for these studies are listed in **Supplemental Table 3**. The data was log transformed in order to permit parametric statistical analyses.

## Murine Plasma Cholesterol profiles and lipoprotein fractions

Total plasma cholesterol levels were determined using an enzymatic colorimetric assay kit (Wako Pure Chemical Industries, Ltd, 439-17501). The lipoprotein cholesterol distribution profile (VLDL, LDL, HDL) was evaluated in pooled plasma samples after fractionation by fast protein liquid chromatography (FPLC) gel filtration on a single Superose 6 column.

## PCSK9 Measurement

Plasma levels of PCSK9, as well as the PCSK9 content of the supernatant of HepG2 cells were measured using an ELISA kit specific to murine PCSK9 (R&D, DY3985) as per manufacturer’s instructions. Samples were diluted 1:100 in 1% BSA and a PCSK9 standard curve was generated for each experiment. PCSK9 levels in the lipid sub-fractions identified using FPLC were also determined using this kit.

## Murine Liver Tissue Western Blotting

Protein was harvested from the livers of male and female mice for SDS-PAGE. Multiple Western blots were obtained for each protein assayed, and representative data are displayed. Mouse liver tissue (200-300 mg) was placed inside a Precellys® homogenizing tube with 4 glass beads and 500 µL of RIPA lysis buffer (50 mM Tris HCL pH 7.4, 1% Triton X-100, 0.5% Sodium Deoxycholate, 150 mM NaCl, 2mM EDTA, protease inhibitor cocktail). Tissue was homogenized during two 30 sec cycles at 6,000 RPM. The lysate was transferred to 2 mL tubes and further processed using a handheld sonicator at 50% amplitude for two 3 sec pulses, centrifuged 45 mins at 16,000*×*g, and the supernatant layer below the fat and above the pellet was transferred to a new tube. Samples were centrifuged again for 10 mins at 16,000*×*g and the supernatants were transferred to new tubes. Protein lysates were quantified using a Bio-Rad DC Protein Assay, with equal amounts of protein (40-80 µg per well) loaded onto 10% Tris-Glycine gels for SDS-PAGE before being transferred to 0.2 μm nitrocellulose membranes ON at 4°C. Membranes were blocked in 5% Milk in TBS-Tween for 1hr, washed, incubated in fresh primary antibodies with 5% BSA ON at 4°C, washed, and incubated in secondary antibody before ECL detection using an ImageQuant LAS4000 imager (GE Healthcare). The primary antibodies used for Western blots with murine liver lysates were: rabbit anti-PCSK9 (1:1,000; ab31762, Abcam), rabbit anti-LDLR (1:5,000; Abcam ab52818), mouse anti-Vinculin (1:2,000; Sigma Aldrich V9131) and mouse anti-*β*-actin (titre: 1:10,000; sc-47778, Santa Cruz Biotechnology). The secondary antibodies used for WB (1:20,000) were: anti-rabbit IgG HRP (Abcam ab97051) and anti-mouse IgG HRP (Jackson Immunoresearch 115-035-062).

## Murine Plasma Glucose & AST

Pooled murine plasma samples from control and treatment groups were sent to Calgary Laboratory Services (Calgary, AB, Canada) for measurement of glucose and aspartate aminotransferase (AST), and the results of these analyses are listed in **Supplemental Table 2**.

## Generation of a Rabbit anti-HSP27 Polyclonal Antibody(PAb)

To study the role of anti-HSP27 antibodies *in vitro* we generated IgG Polyclonal Antibodies (PAb) to rHSP27. A rabbit polyclonal antibody mimicking the human HSP27 autoantibody was produced according to the standard procedure by Cedarlane Laboratories and meeting the requirements of the Canadian Council on Animal Care. Briefly, two rabbits were injected with 0.2 mg of rHSP27 produced in the lab. After 28, 48, 66 days, the rabbits were immunologically boosted using 0.2 mg rHSP27, to increase the quantity of the resulting PAb. Rabbit sera was collected on day 78. The immunization efficiency was determined by indirect ELISA using plates coated with rHSP27. The final serum was loaded to a 5 ml Protein G affinity column (GE Healthcare) for IgG affinity purification. After a 100 ml buffer A (50 mM PBS buffer containing 200 mM NaCl) wash, antibody was eluted by buffer B (20 ml 20 mM sodium acetate, pH = 2.5). The eluted solution was immediately neutralized by buffer C (400 mM PBS, pH = 8.0). The antibody solution was then concentrated using a 15 ml centricon tube with a 30 kDa molecular weight cutoff filter (Millipore Sigma) and the buffer exchanged with buffer A for future usage. Two milligrams of biotinylated HSP27 was then applied to a 1 ml streptavidin affinity column (GE Healthcare) for antigen affinity purification. The antigen conjugated column was washed with 20 ml Buffer A, loaded with PAb from Protein G purification and incubated for 5–10 mins at 4°C, and then washed by 2×20 ml buffer A. The antigen-specific PAb was then eluted with buffer B and immediately neutralized by buffer C to pH ∼7.0. The final purified HSP27 specific PAb was buffer exchanged with Dulbecco’s phosphate-buffered saline buffer and filtered through 0.2 µm filter for future usage.

To verify the fidelity of this PAb we performed epitope mapping on the HSP27 protein using synthesized HSP27 peptides affixed to a membrane (i.e., the spot blot peptide technique) and compared these epitope mapping results to those obtained when human serum was added.^9^ Briefly, a spot blot membrane was constructed with 49 spots representing overlapping 15 amino acid peptide sequences of the full-length HSP27 protein. These peptides were synthesized on a modified cellulose membrane (Kinexus Bioinformatics Corporation, Vancouver, BC). The membrane was soaked ON in 20 mL of PBS containing 1% BSA (pH 7.5), and then washed three times with PBST. A 1:200 (v/v) diluted plasma was added and the membrane was incubated for 2 hrs at RT. After three washes with PBST, the membrane was incubated in a 1:4000 dilution of mouse monoclonal anti-human IgG-HRP (Jackson Immunoresearch) for 1 hr. After three additional washes with PBST, the interacting peptide dots were developed with an ECL kit (GE Healthcare) for 3–5 mins according to manufacturer’s instructions and imaged by the FluorChem® Q Imaging System (Cell Biosciences, Santa Clara, CA).

Specific 15-mers of the linearized protein reacted with the PAb in a pattern that was identical to that of the naturally occurring AAbs – whether they were derived from the blood of CON or CVD subjects (**Supplemental Fig. 4A**). The binding of the PAb to these epitopes was abrogated by the addition of rHSP27 (100 μg/ml vs. an irrelevant anti-human IgG negative control), thereby indicating that epitopes in the full-length rHSP27 were similar to those on the spot blot (**Supplemental Fig. 4B**). A summary of the HSP27 epitopes is schematically outlined in **Supplemental Fig. 4C**. When the rabbit PAb was applied to the Spot blot the identified epitopes were identical to those note with human sera from CON and CVD subjects, thereby confirming the specificity of the PAb (**Supplemental Fig. 4D**). Finally, we compared the immunoreactivity of this rabbit PAb *vs*. a commercial (goat) anti-HSP27 antibody using Western blotting of rHSP27 and rHSP25 (**Supplemental Fig. 4E**). The addition of excess rHSP27 essentially eliminated an interaction signal for either antibody. In the absence of exogenous rHSP27 the band recognition pattern for various (loaded) amounts of rHSP27 (10 ng, 100 ng, 1 μg) or rHSP25 (100 ng, 1μg) were very similar for these two antibodies.

## *In Vitro* Experiments with Human Hepatocyte Cell Lines

Human hepatic cells (HepG2, ATCC HB-8065) were cultured in MEMα (supplemented with 10% FBS and 1% Pen-Strep) at 37°C in a 5% CO2 humidified environment. For qPCR and Western blot experiments, cells were seeded ON in complete media at a density of 0.5*×*10^6^ cells/well in 6-well plates. The media was replaced with serum-free fresh media and treated as described in each experiment. Cells were harvested for gene expression by qPCR (see **Supplemental Table 3** for primers). Huh7 human hepatocytes, a kind gift from Dr. Thomas Lagace (University of Ottawa Heart Institute, ON, Canada)^10^, were used for immunofluorescent labeling experiments as these cells grow in flat monolayers *in vitro*; whereas HepG2 cells tend to grow in clumps, making visualization of individual cells suboptimal.

## Stable isotope labelling of amino acids in cell culture (SILAC) and Mass Spectrometry

HepG2 cells were grown for 24 hrs in dialyzed FBS and Pen-Strep with either: i) ‘light’ media RPMI 1640 (control cells) or ii) ‘heavy’ media SILAC RPMI 1640 containing heavy isotopes (^13^C_6_^15^N-L-lysine-2HCl and ^13^C_6_^15^N_4_-L-arginine-HCl) with 1 µg/mL rHSP27 plus 5 µg/mL PAb. Protein extracts were obtained after washing the cells with PBS then lysed in a modified RIPA buffer (50 mM Tris-HCl, pH 7.8, 150 mM NaCl, 1% (v/v) NP-40, 0.25% sodium deoxycholate and 1 mM EDTA) with Protease Inhibitor Cocktail (Roche). Lysates were mixed in a 1:1 ratio, reduced with DTT, alkylated, and digested ON with Trypsin. Peptides were separated on an Ultimate 3000 HPLC system (ThermoFisher) coupled to a QTRAP 4500 LC-MS/MS (Sciex). Selected protein target peptides were measured by MRM and data were analyzed and quantitated using Skyline software (MacCoss Lab, University of Washington, Seattle, WA, USA) by generating a ratio of light (PBS control) to heavy (rHSP27 + PAb) for each peptide. Ratios for each protein target were normalized to the housekeeping protein GAPDH and converted to represent the amount of target protein in each treatment relative to control cells.

## Effect of [rHSP27 + PAb] on HepG2 Expression PCSK9, LDLR and HNF1*α* Genes

As per the description of the methodology for the Murine Liver Tissue Q-PCR, RNA was isolated from HepG2 cells and used for Real-time PCR analysis of LDLR, PCSK9, and HNF1*α* gene expression. Prior to RNA harvesting, HepG2 cells were treated with rHSP27 (1 µg/mL) plus PAb (5 µg/mL) or control treatments [PBS, PAb alone (5 µg/mL), rC1 (1 µg/mL) plus PAb (5 µg/mL), and rHSP27 (1 µg/mL) plus a non-specific rabbit polyclonal IgG (5 µg/mL; Jackson ImmunoResearch 304-005-003). The specific human primer sequence pairs, including those for the control gene, hypoxanthine-phosphoribosyltransferase (hHPRT1), are provided in **Supplemental Table 3**. RNA expression is presented as log fold change.

## Changes in Intracellular Cholesterol Concentration and LDLR Expression in HepG2 Cells

HepG2 cells were grown in 6 well plates containing 0.5-1.0*×*10^6^ cells. The cell culture media was then supplemented with either 10μg/ml LDL or not. Intracellular cholesterol levels were determined using a method disseminated by the laboratory of Dr. Alan Daugherty.^11^

## LDLR and PCSK9 Promoter Activity Assays

PCSK9 and LDLR promoter firefly luciferase reporter constructs, kindly provided by Dr. Jingwen Liu (Palo Alto Veterans Institute for Research, California, USA), were co-transfected with a constitutively expressed *Renilla* luciferase vector as a transfection control, and the dual luciferase activities were measured. The PCSK9 promoter construct was incorporated in the plasmids pGL3-PCSK9-D1 (referred to as PCSK9-D1). For the LDLR promoter we used the previously described constructs: full length (pLDLR-1192) and core (pLDLR-234 containing only the core regulatory elements SRE-1 and SP1 sites) ^12, 13^. The plasmid pBLSV40 containing *Renilla* luciferase under the control of the constitutive SV40 promoter was generously donated by Dr. David Proud (University of Calgary, Calgary, AB). HepG2 cells were grown in complete media (with 10% FBS and antibiotics) and seeded ON at 4*×*10^4^ cells/well in a 48-well culture plates then transiently transfected with 500 ng of the PCSK9-D1 or LDLR-234 promoter constructs and 25 ng of pBLSV40 using 1 uL of Lipofectamine®2000 Transfection Reagent (ThermoFisher). Cells were then treated with recombinant proteins (rHSP27, rC1), antibodies (unrelated IgG control, PAb), and combinations thereof for 24 hrs. PCSK9 or LDLR promoter-driven firefly luciferase activity and control renilla luciferase activity were measured using the Firefly & Renilla Luciferase Single Tube Assay Kit (#30081-1; Biotium, Fremont, CA) according to the manufacturer’s protocol, with some minor modifications. Briefly, cells were washed with PBS then lysed in 110 uL of passive lysis buffer. Firefly luciferase activity was measured first by combining 30 uL of cell lysate with 50 uL of firefly working solution in a 12*×*75mm round-bottom tube, measuring luminescence, then adding 5 uL of *Renilla* working solution to the same tube and again measuring luminescence. Luminescence was detected using a Monolight 3010 Luminometer (BD Biosciences). Absolute firefly luciferase activity was normalized against *Renilla* luciferase activity to correct for transfection efficiency. Triplicate wells were assayed for each transfection condition, and 3 independent transfection assays were performed for each reporter construct. The data was log transformed in order to facilitate parametric statistical analyses.

## Effect of HSP27 on Extracellular PCSK9 Levels

Two experiments were performed to assess the effect of HSP27 on PCSK9 secretion in HepG2 cells, each involving a manipulation:

### i) Knockdown of HSP27 with shRNA

To assess the effect of HSP27 on PCSK9 secretion in HepG2 cells the expression of HSP27 was knocked down with small hair pin RNA (shRNA). shRNA sequences were designed and subcloned into the pRNAi-H1-Hyg plasmid (Biosettia, San Diego, CA) in order to attenuate the expression of HSP27 in HepG2 cells. Using electroporation, plasmids were transfected into HepG2 cells and stable cell lines were selected using hygromycin. Cells treated with no plasmid or a scrambled RNA sequence were used as controls. To confirm that HSP27 protein was reduced with this treatment, cells were lysed and treated with 1% benzonase nuclease (Sigma Aldrich E1014), and the protein extract was stored in Gel Loading Buffer [3% SDS (W/V), 10% Glycerol, 500 mM Tris-HCl pH6.8, 1% ß-mercapto-ethanol, 0.05% Bromophenol blue] at −20°C. Western blotting was performed using the following primary antibodies: the rabbit polyclonal anti-HSP27 generated by our lab (PAb, 1:5000) and a mouse anti-β-actin antibody (1:5000, Abcam ab184220). The reductions in HSP27 protein in the cells treated with HSP27 shRNA (as quantified using densitometric analysis) were 66% and 71% compared to no treatment and Scramble shRNA treatment, respectively. The secondary antibodies used for WB (1:20,000) were: anti-mouse IgG HRP (Jackson Immunoresearch 115-035-062) and anti-rabbit IgG HRP (Abcam ab97051). To establish the effect of knocking down HSP27 expression on extracellular PCSK9 levels, the aforementioned cells were grown in culture and the abundance of PCSK9 in the extracellular fluid (cell supernatant) was assessed using an ELISA for human PCSK9 (R&D DY3888).

### ii) Increased PCSK9 secretion with statin treatment

Statin-induced PCSK9 secretion was assessed in the culture media of HepG2 cells treated ON (16 hrs) with Atorvastatin (AV; 1 µM and 10 µM, Millipore Sigma 189291) without or with rHSP27 (1 µg/mL) plus PAb (5 µg/mL) using the aforementioned human PCSK9 ELISA.

### Immunofluorescence staining of NF-Kb p65

For immunostaining, rabbit polyclonal affinity purified antibody to p65 (sc-109) (Santa Cruz Biotechnology) was used. 10% normal goat serum (Sigma Aldrich) was used as a blocking buffer, eliminated all non-specific binding of the secondary antibody. Huh7 human hepatocyte cells were cultured on glass coverslips and fixed with 4% paraformaldehyde (at least 1hr, RT), washed with PBS. We did not use HepG2 cells for this experiment, as they tend to grow in clumps, whereas Huh7 cells grow in flat monolayers *in vitro*. Huh7 cells were treated with PAb (5 μg/ml), rHSP27 (1 μg/ml) and [rHSP27+PAb] for 45 minutes. All reagents were diluted in PBS, and coverslips were immersion washed in PBS between each staining step. Coverslips were incubated sequentially with 0.5% Triton-X100 (Sigma) (10 mins, RT), blocking buffer 10% normal horse serum (30 mins, RT), primary antibody to the NF-*κ*B P65 subunit (rabbit anti human BF-kB p65 antibody from Santa Cruz Biotechnology, Inc., cat# sc-109; 1:60; Dallas, Texas, USA) diluted in PBS (ON, 4°C) and secondary antibody diluted in PBS (30 mins, RT). Nuclei were counterstained with Hoechst dye (Pierce Biotechnology, Rockford, IL) for 8 mins, then wash with PBS. Coverslips were mounted on to glass slides using mounting media 40% Gly (Vector). Photomicrographs were obtained using a fluorescent microscopy.

### LDLR Expression and NF-*κ*B Pathway Blockade

As we previously showed that HSP27 activates the NF-*κ*B pathway via TLR4 to enact the transcription of various genes that are relevant to atherosclerosis (both pro- and anti-inflammatory factors),^1, 6^ we sought to determine if this pathway was involved in the upregulation of LDLR expression. HepG2 cells were seeded in 24 well plates and treated for up to 24 hrs with Control (DMSO vehicle), the NF-*κ*B inhibitor BAY 11-7082 (10μM), [rHSP27 (1μg/ml) + PAb (5μg/ml)] or BAY 11-7082 (10μM) plus [rHSP27 (1μg/ml) + PAb (5μg/ml)]. RNA and protein were harvested. qPCR was performed using the housekeeping gene hHPRT1 to normalize results before log transforming the data. Western blotting was completed using a polyclonal rabbit anti-human LDLR antibody (1:2,000; Biovision, Milpitas, CA) at RT for 2.5 hrs, followed by a donkey anti-goat antibody (1:7,500; Santa Cruz, sc-2020) for 1 hr. The LDLR protein band densities were normalized to those of the housekeeping protein, Vinculin.

### Duolink Proximity Ligation Assay

The Duolink proximity ligation assay (PLA) was used to identify the interaction between rHSP27 and the PAb in HuH7 hepatocytes. Cells were plated in 24-well plate with cover slides (12 mm diameter) and cultured until they were approximately 70% confluent before being placed in fresh FBS-free media for 6 hrs then treated with rHSP27 (1ug/ml) or rHSP27 (1ug/ml) + PAb (5ug/ml) for 24 hrs. The slides were then rinsed thrice with PBS and fixed with 10% neutral formalin (Sigma) for 10 mins and permeabilized with 0.1% Triton X-100 in PBS for 15 mins at RT. Thereafter, the cells were blocked with 3% goat serum in PBS at 37°C for 1hr then incubated with blocking buffer (Olink Bioscience, Uppsala, Sweden) ON at 4°C before incubating with a primary goat anti-rabbit antibody at 37°C for 1hr. The PLA probe anti-goat “minus” and the PLA probe anti-rabbit “plus” were then added for another hr at 37°C. For additional negative controls, slides were treated with PBS, PAb (alone), rC1 or [rC1 + PAb].

Proximity ligation was then conducted *in situ* according to the manufacturer’s instructions (Olink Bioscience). If the PLA probes are in close proximity, bound antibody-oligonucleotide conjugates will ligate for 30 mins at 37°C. After washing, slides were incubated with amplification solution, and ligated templates were amplified for 100 mins at 37°C. Slides were then washed twice with wash buffer for 10 mins each and mounted with DPI. Microscopic images were obtained with an Olympus (Center Valley, PA) Fluoview FV10i Confocal Microscope at 200*×*, with the selected fluorescent dye of Texas Red (excitation 598 nm and emission 634nm). This technique generates a fluorescent signal when rHSP27 and the PAb co-localize within 40 nm of each other.

### Statistical Analyses for the non-clinical data

Continuous variables are expressed as mean ± standard error of the mean (SEM) and the comparison of unpaired treatment groups was performed using a Student’s t-test. Quantitative PCR gene expression data were log-transformed and are reported as log-fold change in order to more normally distribute the data and permit parametric statistical analyses. All analyses were performed in a blinded fashion using GraphPad Prism 8 (GraphPad Software, La Jolla, CA). A two-tailed α level of 0.05 was used to define statistical significance.

---

**Supplemental Fig. 1:**
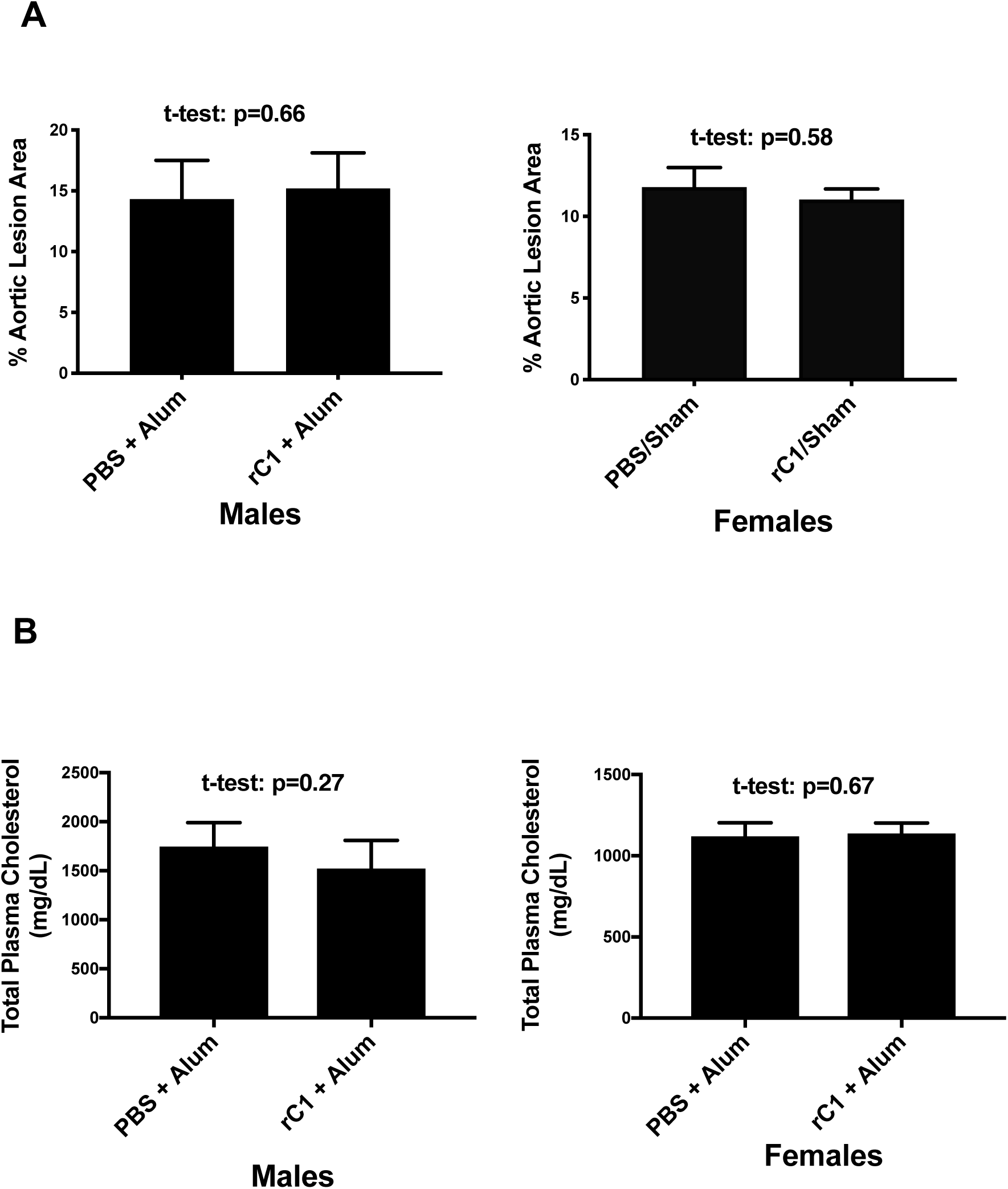
Validation of rC1 Control Treatment As rC1 was generated in the same manner as rHSP25 using *E. coli*, it is the ideal comparator for rHSP25 active treatments as it can balance any unforeseen confounding factors, such as the potential for minute levels of endotoxin contamination from the preparation of the recombinant proteins. Hence, we compared vaccination with [rC1 + Alum] *vs.* [PBS + Alum] in male and female mice, looking specifically at atherosclerotic burden and plasma cholesterol levels. A) There is no difference in aortic lesion area between [PBS + Alum] *vs.* [rC1 + Alum] vaccination groups, with the male data on the left and the female data on the right. B) There was no difference in the total plasma cholesterol levels between [PBS + Alum] *vs*. [rC1 + Alum] vaccination groups, with the male data on the left and the female data on the right.

**Supplemental Fig. 2:**
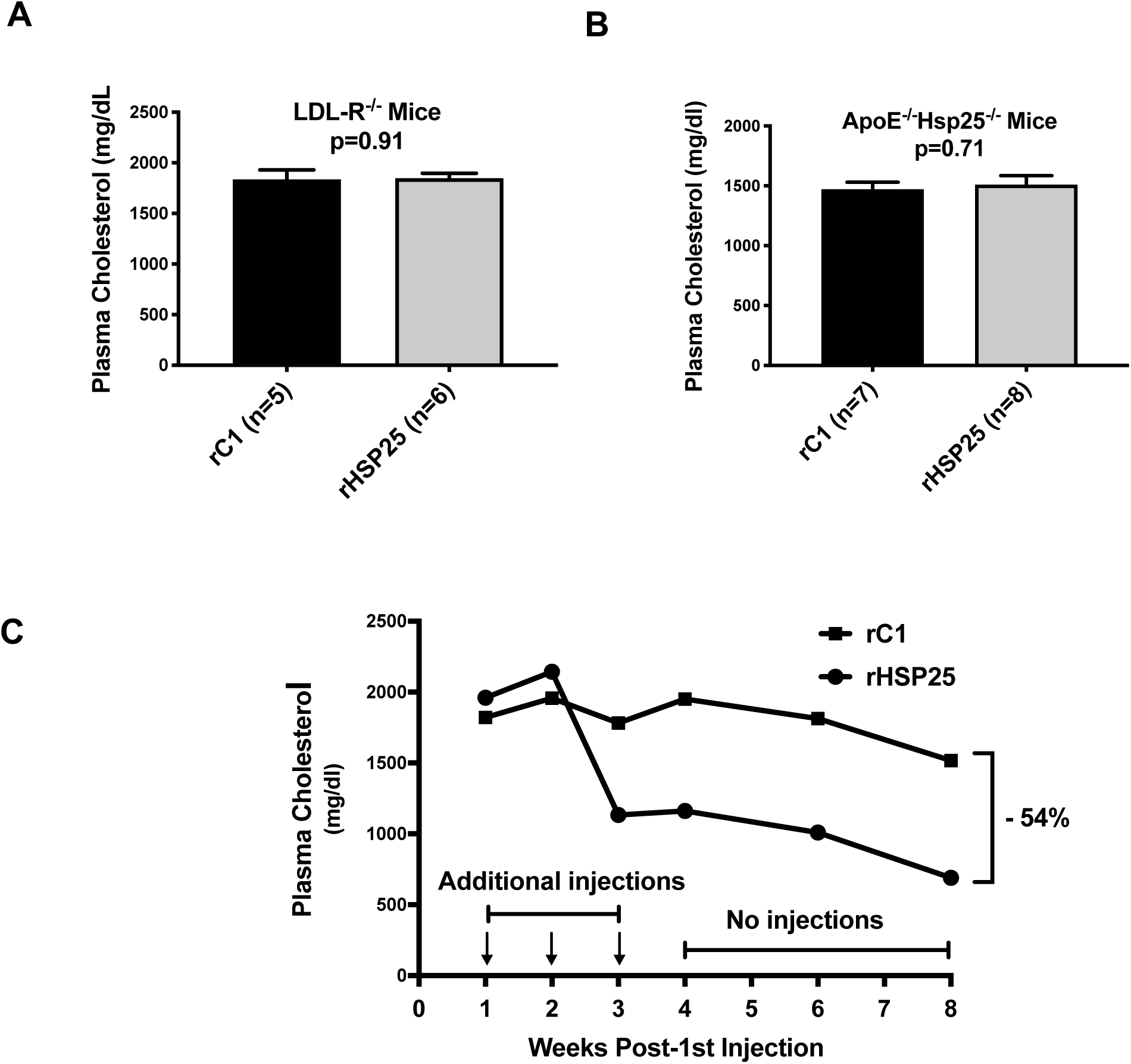
rHSP25 Vaccination Experiments A) Plasma cholesterol levels in *LDL-R^-/-^* mice were unaltered after vaccination with rHSP25 (*vs.* rC1), suggesting that LDL-R is required for rHSP25 vaccination to successfully attenuate atherogenesis. B) Plasma cholesterol levels in *ApoE^-/-^HSP25^-/-^* mice were unchanged after vaccination with rHSP25 (*vs.* rC1), suggesting that basal levels of HSP25 are required for rHSP25 vaccination to successfully attenuate atherogenesis. C) Plasma cholesterol levels remain markedly lower 5 weeks after the last rHSP25 vaccination in *ApoE^-/-^* mice.

**Supplemental Fig. 3:**
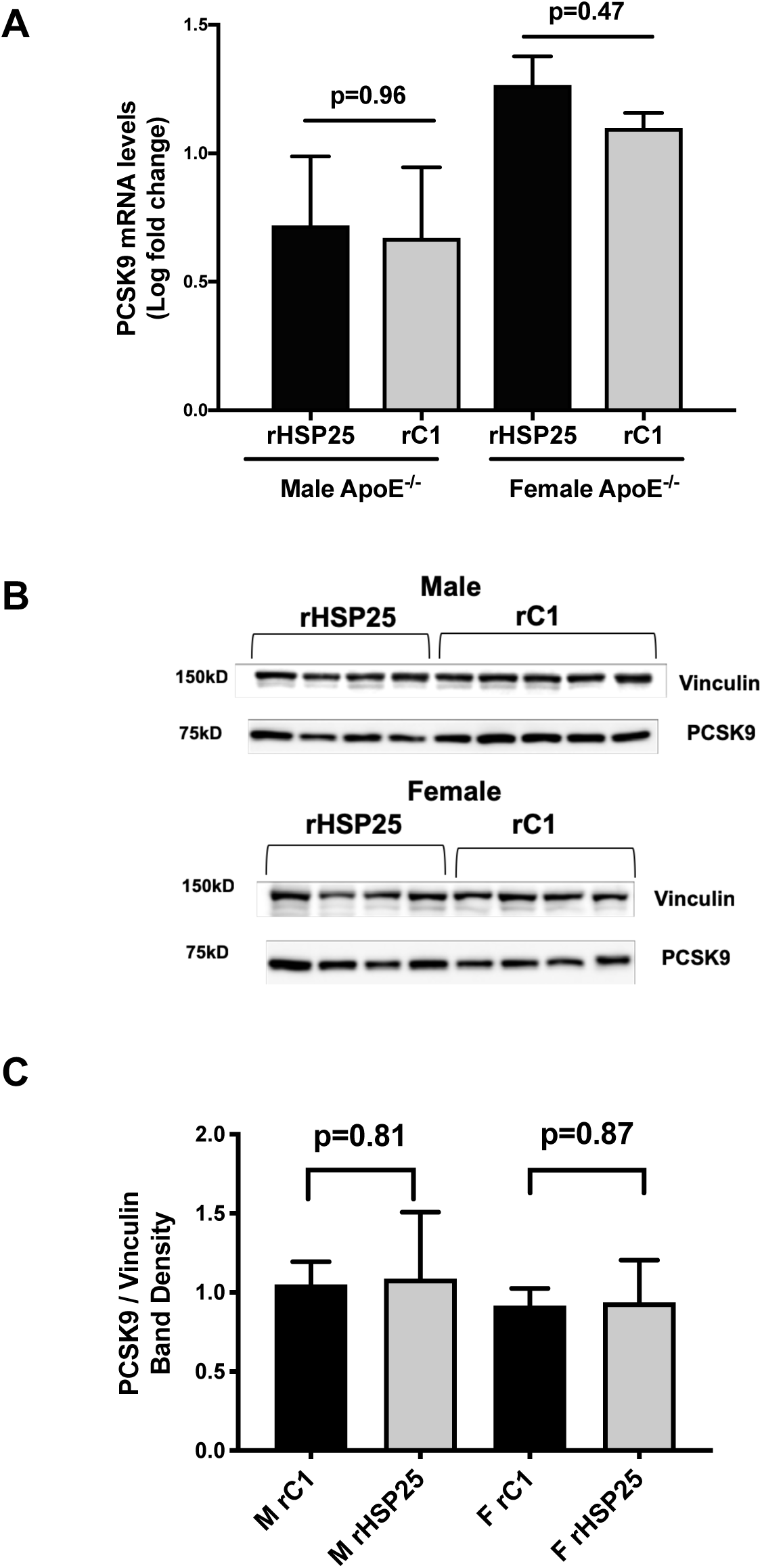
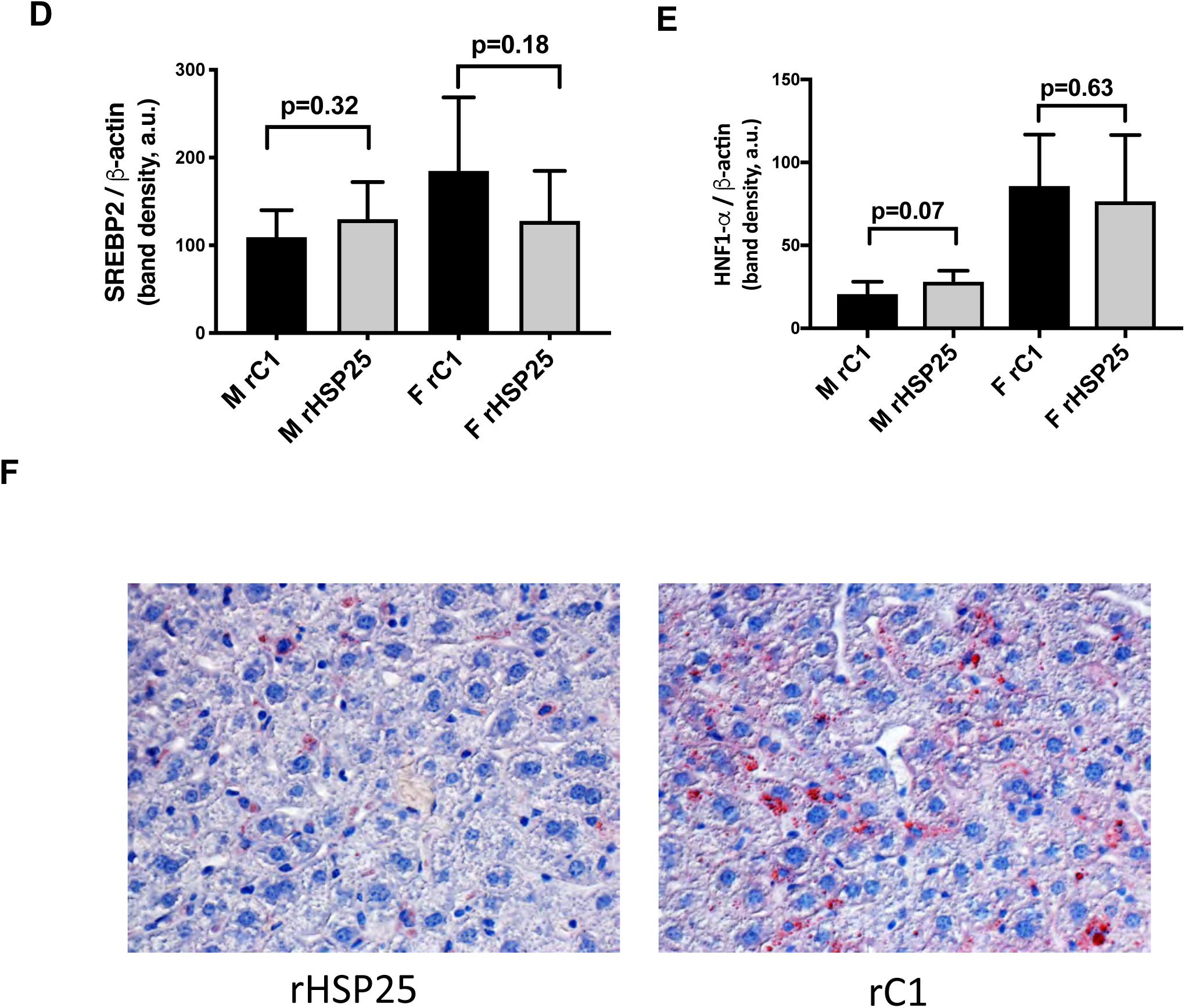
PCSK9 data A) PCSK9 mRNA expression is similar in the male and female mice vaccinated with rC1 *vs*. rHSP25. B) & C) PCSK9 protein expression is similar in the male and female mice vaccinated with rC1 *vs.* rHSP25. Western blot showing PCSK9 expression. Vinculin was used as a housekeeping protein. PCSK9 band density was normalized to that of vinculin. D) and E) Western blotting revealed that SREBP2 and HNF1*α* protein expression was unaltered with rHSP25 vaccination compared to rC1. *β*-actin was used as the housekeeping protein for normalization of the target protein bands. F) Hepatic Oil Red O content (original photo: *×*400 magnification)

**Supplemental Fig. 4:**
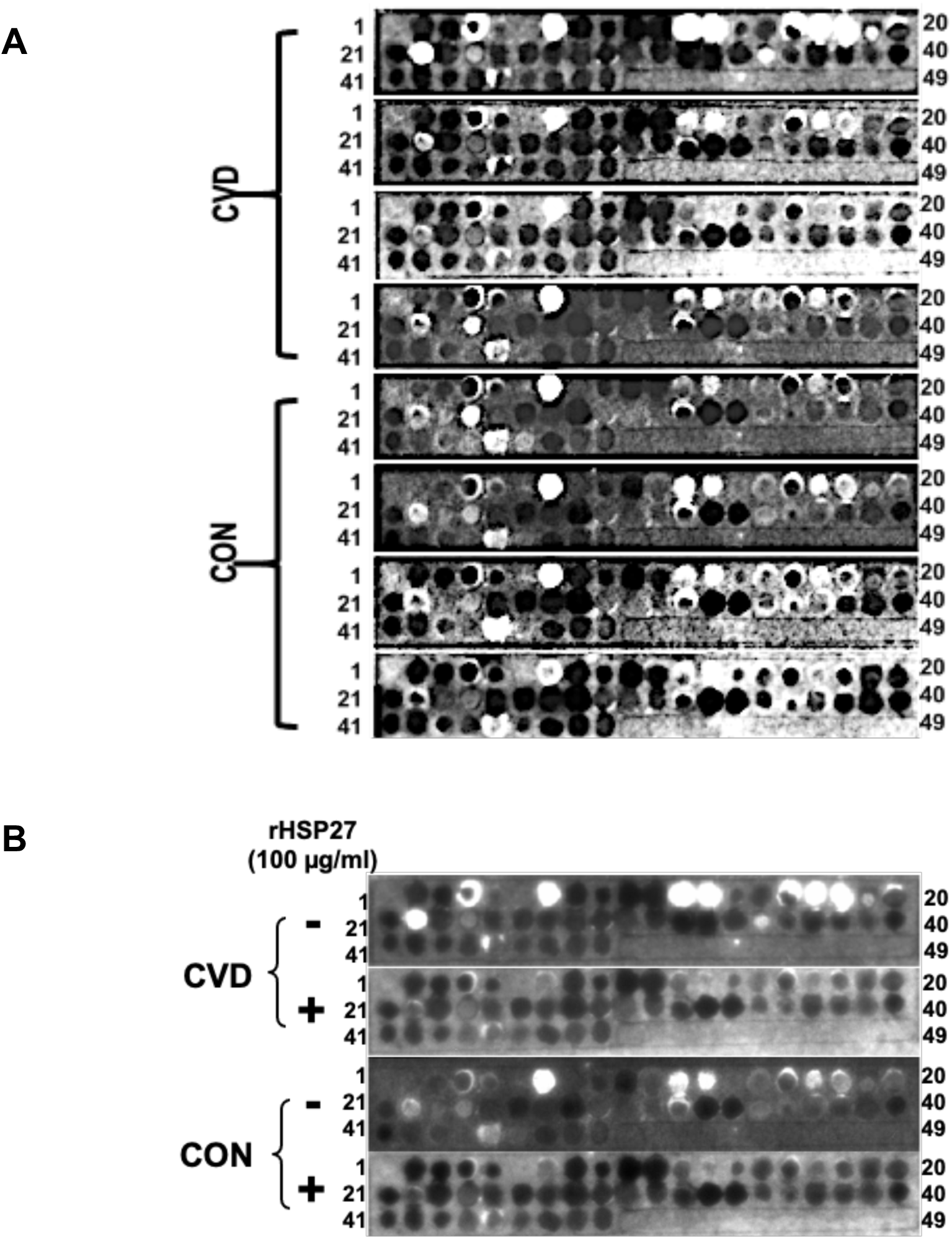
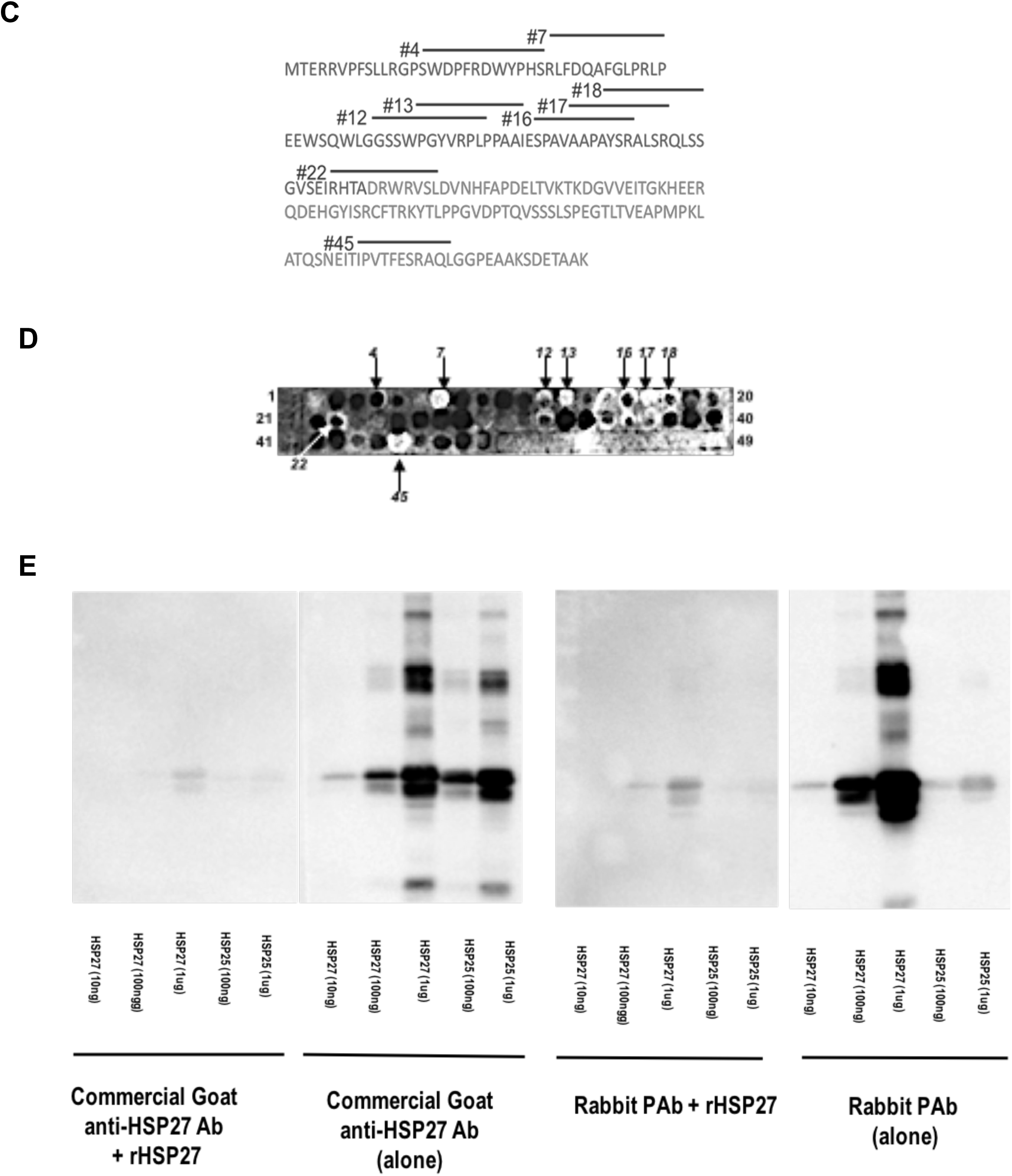
HSP27 Epitope Mapping and Validation of anti-HSP27 Polyclonal IgG Antibody A) A Spot Blot was constructed with 49 dots representing overlapping 15-amino acid sequences of the HSP27 protein sequence. The interaction of an antibody with a specific dot is indicated by the white chemiluminescence signal. Natural anti-HSP27 antibodies from the blood cardiovascular disease patients (CVD; n=4) and health controls (CON; n=4) identified the same dots, with no major differences between CVD and CON subjects. B) Adding rHSP27 displaced natural anti-HSP27 antibodies from CVD and CON subjects, thereby highlighting the specificity of the Spot Blot interaction. C) Mapping HSP27 autoantibody interaction epitopes (numbered heavy lines above amino acid sequence). Most epitopes are in the N-terminus of HSP27. D) The rabbit anti-HSP27 IgG Polyclonal (PAb) interacts with the Spot Blot in a patterns similar to that of the human blood samples shown in A). E) Western blotting using a commercial goat anti-HSP27 antibody (left sided panels) or the generated rabbit PAb (right panels). For each antibody the individual left and right panels shows the blot results with or without exogenous rHSP27 (to block the interaction). Hence, the pattern of protein identification with these two antibodies are very similar.

**Supplemental Fig. 5:**
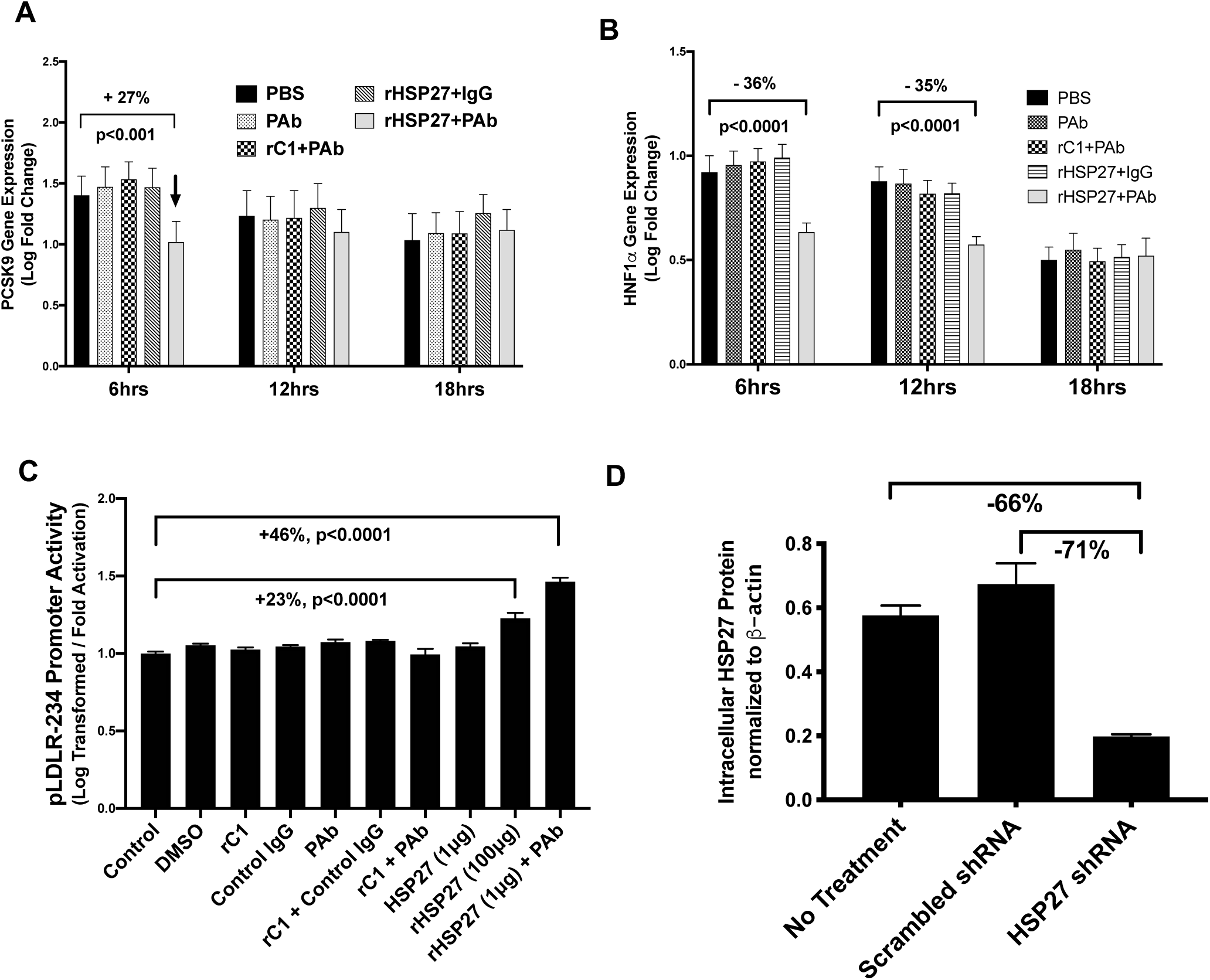
PCSK9 and LDLR data A) - B) Time course of changes in gene expression assessed for PCSK9 and HNF1*α* post-treatment with [rHSP27 (1 µg/mL) plus PAb (5 µg/mL)] or control treatments [PBS, PAb alone (5 µg/mL), rC1 (1 µg/mL) plus PAb (5 µg/mL), and rHSP27 (1 µg/mL) plus non-specific polyclonal IgG (5 µg/mL)]. Changes in gene expression were assessed by qPCR and expressed as log fold change. Relative to control treatments, [rHSP27 + PAb] decreased both PCSK9 and HNF1*α* expression, although these changes were relatively modest, and for PCSK9 transient. C) HepG2 cells transfected with the core LDLR promoter construct containing only the Sterol Response Element-1 (SRE-1) and Sp1 regulatory element sites from −142 to +35, that employs firefly luciferase to demonstrate promoter activation. Various controls and treatments are outlined, with the results expressed as log transformed activation. Compared to control LDLR promoter activity strongly increased with [rHSP27 + PAb] treatment. D) Small hair pin RNA (shRNA) sequences designed to knockdown the expression of HSP27. Treatment with HSP27 shRNA reduced the expression of HSP27 (as measured by western blot densitometry) by 66% and 71% relative to no treatment and scrambled shRNA.

**Supplemental Fig. 6:**
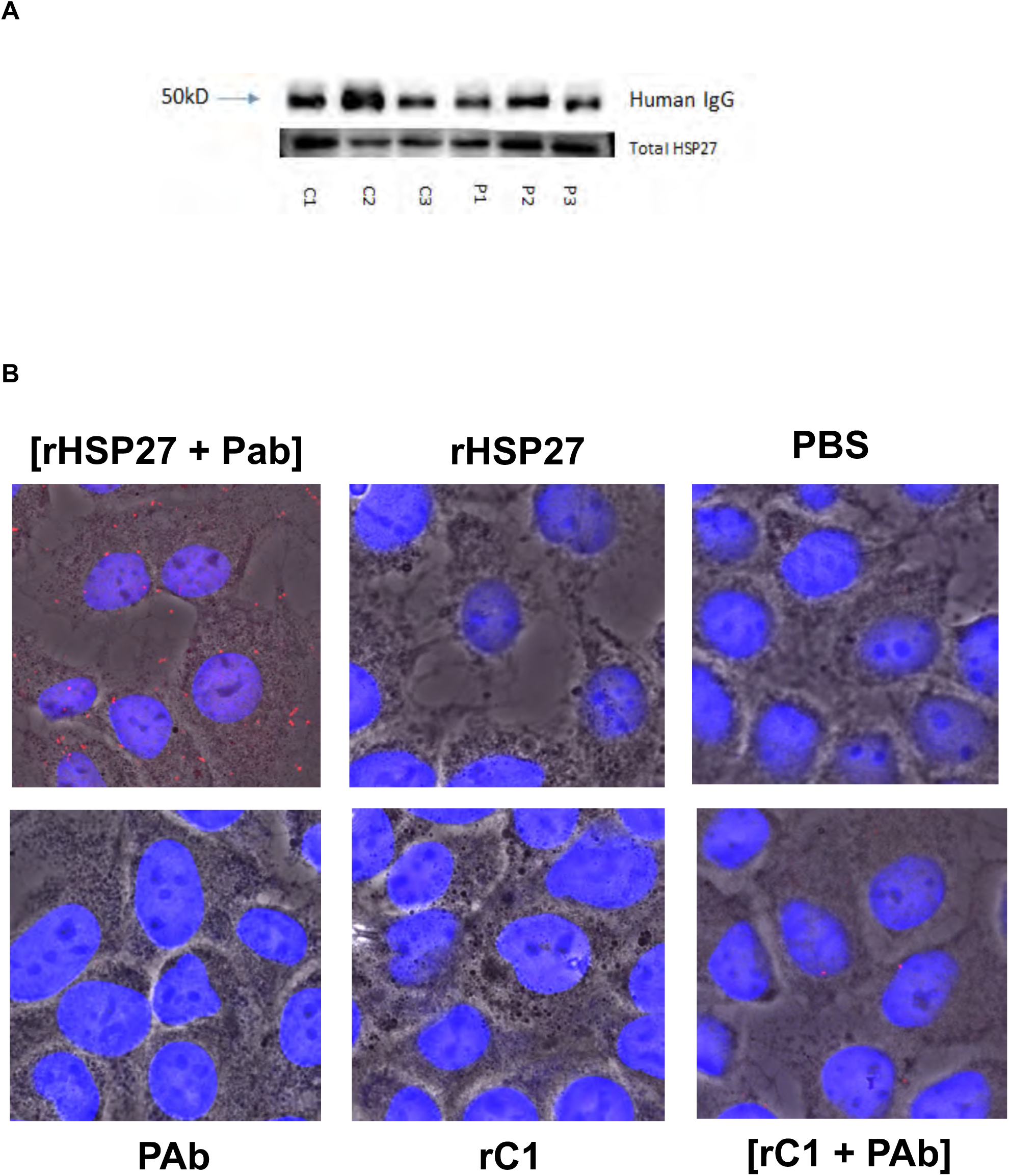
Demonstration of HSP27 Immune Complex A) Components of HSP27 IC detected in pooled human blood samples. Ultracentrifugation of blood was used to obtain a pellet that contained microvesicles that were resolved using SDS-PAGE followed by Western Blotting. Bands show immunodetection of HSP27 and human IgG from both health controls (C1-3) and CVD patients (P1-3), with 6 subjects per group. B) Photomicrographs of the Duolink® Proximity Ligation Assay (PLA), a highly specific and sensitive means of demonstrating *in situ* the formation of a HSP27 IC. A red fluorescent signal is generated when rHSP27 and the PAb co-localize within 40 nm of each other. In the top left panel, the red fluorescent signal documents the presence of HSP27 IC in hepatocytes treated with rHSP27 and PAb (200*×* magnification). All other panels show control experiments and the absence of a specific interaction signal.

**Supplemental Table 1:**
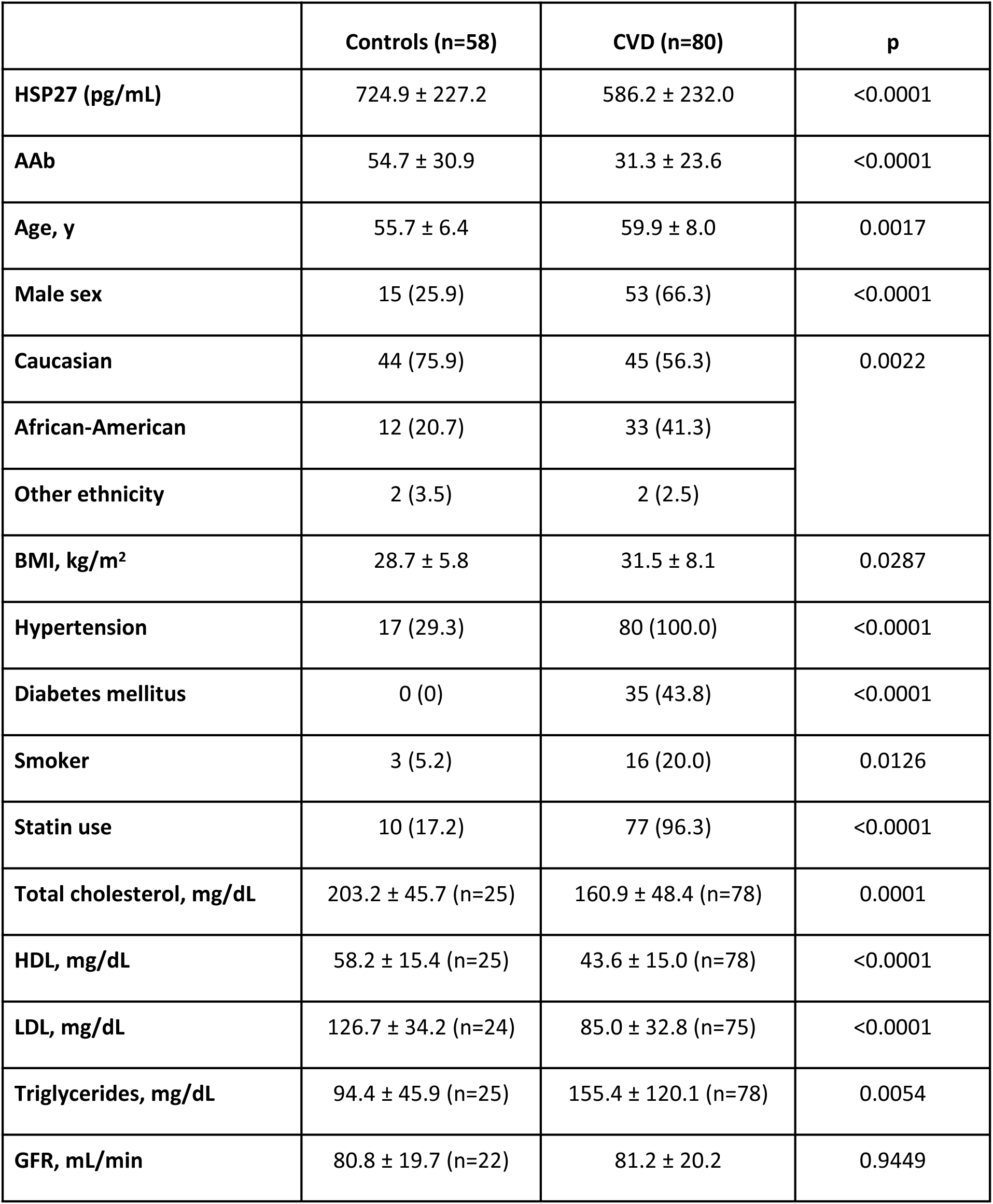
Characterization of the NIH study cohorts: healthy controls *vs.* CVD patients. The following abbreviations were used: HSP27: heat shock protein 27; AAb: antibodies to HSP27; CVD = cardiovascular disease cohort; CON = healthy control cohort; BMI = body mass index; GFR = glomerular filtration rate; HDL = high density lipoprotein; LDL = low density lipoprotein.

|  | **Controls (n=58)** | **CVD (n=80)** | **p** |
| --- | --- | --- | --- |
| **HSP27 (pg/mL)** | 724.9 ± 227.2 | 586.2 ± 232.0 | <0.0001 |
| **AAb** | 54.7 ± 30.9 | 31.3 ± 23.6 | <0.0001 |
| **Age, y** | 55.7 ± 6.4 | 59.9 ± 8.0 | 0.0017 |
| **Male sex** | 15 (25.9) | 53 (66.3) | <0.0001 |
| **Caucasian** | 44 (75.9) | 45 (56.3) | 0.0022 |
| **African-American** | 12 (20.7) | 33 (41.3) |  |
| **Other ethnicity** | 2 (3.5) | 2 (2.5) |  |
| **BMI, kg/m<sup>2</sup>** | 28.7 ± 5.8 | 31.5 ± 8.1 | 0.0287 |
| **Hypertension** | 17 (29.3) | 80 (100.0) | <0.0001 |
| **Diabetes mellitus** | 0 (0) | 35 (43.8) | <0.0001 |
| **Smoker** | 3 (5.2) | 16 (20.0) | 0.0126 |
| **Statin use** | 10 (17.2) | 77 (96.3) | <0.0001 |
| **Total cholesterol, mg/dL** | 203.2 ± 45.7 (n=25) | 160.9 ± 48.4 (n=78) | 0.0001 |
| **HDL, mg/dL** | 58.2 ± 15.4 (n=25) | 43.6 ± 15.0 (n=78) | <0.0001 |
| **LDL, mg/dL** | 126.7 ± 34.2 (n=24) | 85.0 ± 32.8 (n=75) | <0.0001 |
| **Triglycerides, mg/dL** | 94.4 ± 45.9 (n=25) | 155.4 ± 120.1 (n=78) | 0.0054 |
| **GFR, mL/min** | 80.8 ± 19.7 (n=22) | 81.2 ± 20.2 | 0.9449 |

**Supplemental Table 2:** Mean glucose and aspartate transaminase (AST) levels for each of the murine cohorts at the end of the study, with no apparent differences between the rHSP25 and rC1 vaccination groups.

| Treatment / Mice | Glucose (mmol/L) | AST (U/L) |
| --- | --- | --- |
| rHSP25 / Male | 13.7 | 71 |
| rC1 / Male | 12.9 | 71 |
| rHSP25 / Female | 12.3 | 129 |
| rC1 / Female | 11.2 | 165 |

**Supplemental Table 3:**
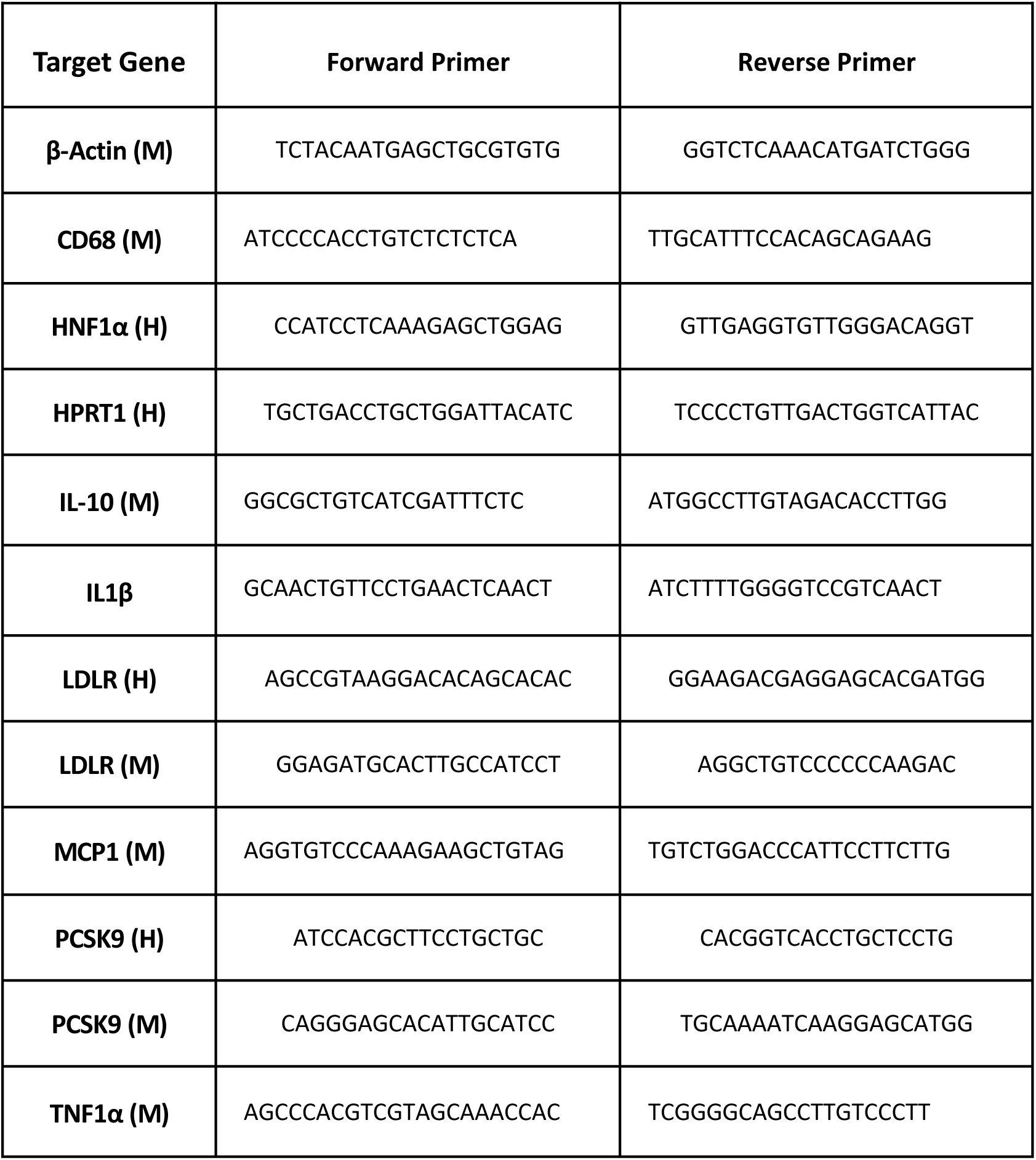
Sequences of the qPCR primers used to assess expression of genes in mice (M) and Humans (H)

